# Patterns of African and Asian admixture in the Afrikaner population of South Africa

**DOI:** 10.1101/542761

**Authors:** N Hollfelder, JC Erasmus, R Hammaren, M Vicente, M Jakobsson, JM Greeff, CM Schlebusch

## Abstract

The Afrikaner population of South Africa are the descendants of European colonists who started to colonize the Cape of Good Hope in the 1600’s. In the early days of the colony, mixed unions between European males and non-European females gave rise to admixed children who later became incorporated into either the Afrikaner or the “Coloured” populations of South Africa. Ancestry, social class, culture, sex ratio and geographic structure affected admixture patterns and caused different ancestry and admixture patterns in Afrikaner and Coloured populations. The Afrikaner population has a predominant European composition, whereas the Coloured population has more diverse ancestries. Genealogical records estimated the non-European contributions into the Afrikaners to 5.5%-7.2%. To investigate the genetic ancestry of the Afrikaner population today (11-13 generations after initial colonization) we genotyped ~5 million genome-wide markers in 77 Afrikaner individuals and compared their genotypes to populations across the world to determine parental source populations and admixture proportions. We found that the majority of Afrikaner ancestry (average 95.3%) came from European populations (specifically northwestern European populations), but that almost all Afrikaners had admixture from non-Europeans. The non-European admixture originated mostly from people who were brought to South Africa as slaves and, to a lesser extent, from local Khoe-San groups. Furthermore, despite a potentially small founding population, there is no sign of a recent bottleneck in the Afrikaner compared to other European populations. Admixture among diverse groups during early colonial times might have counterbalanced the effects of a founding population with a small census size.

**SIGNIFICANCE STATEMENT:** Afrikaners are a southern African ethnic group primarily descended from colonial settlers (population ~2.8–3.5 million). Genome-wide studies might offer interesting insights into their ancestry, not the least due to South Africa’s history of segregationist laws known as “apartheid”, resulting in an expectation of low levels of admixture with other groups. Originating from a small founder population, their genetic diversity is also interesting. In our genome-wide study of 77 Afrikaners we found their majority ancestry (average 95.3%) came from Europeans, but almost all Afrikaners had admixture from non-Europeans (Africans and Asians). Despite their small founding population, we found no signs of decreased genetic diversity. Admixture among diverse groups during colonial times might have counterbalanced effects of a small founding population.

## INTRODUCTION

The seventeenth century European colonization of the southern tip of Africa resulted in the influx of two groups of people, European colonists and slaves. The subsequent admixture between these external groups and the local southern African Khoe-San populations resulted in two admixed populations – the Afrikaner population and the “Coloured” population of South Africa (1). (In this article we use the term “Coloured” following the current-day continued use of the term as self-identification (2)).

While both the Afrikaner and Coloured populations have ancestry from many populations from different continents, the ancestry proportions differ substantially between the groups. The admixture proportions of these populations do not reflect the historical local census sizes of the parental populations (supplementary text), rather, ancestry, social class, culture, sex ratio and geographic structure affected admixture patterns (3-7).

The most dominant contribution to the Afrikaner population came from European immigrants ((8-10) and supplementary text), whereas the Coloured population has more diverse ancestries (11-20). The colonization of southern Africa started in 1652 when Jan van Riebeeck established a refreshment station for the Dutch East India Company (DEIC) at the Cape of Good Hope (Cape Town today). In 1657 a few employees of the DEIC were released from their services to start farming (4) (numbering 142 adults and children in 1658). They were initially called free citizens and later Afrikaners (supplementary text). The DEIC continued to support the settlement of Dutch nationals and other Europeans by offering free passage to the Cape and granting land for farming (4). The other major sources of immigrants were 156 French Huguenots that arrived in 1688 and an unknown number of German laborers and soldiers that were financially marginalized (4). Estimates vary but Dutch, German and French respectively contributed 34%-37%, 27%-34% and 13%-26% ((8-10) and supplementary text). Immigrants from Europe continued to arrive at the Cape from 1658 onwards, but the very rapid population growth of the local established population (almost 3% per annum; (5, 21) and supplementary text) and lower fecundity in the new immigrants (5) meant that the genetic contribution of later arrivals was less ((8) and supplementary text).

While the DEIC did not encourage admixture with local populations and slaves, the strongly male-biased ratio of immigrants lead to mixed-ancestry unions (3), especially between European males and non-European females (however there are recorded cases of European women marrying non-European men) (7). The offspring from these unions were frequently absorbed into the Afrikaner population (8). As time progressed relationships between Europeans and non-Europeans became more infrequent (8) and as early as 1685, commissioner van Rheede outlawed marriages between Europeans and non-Europeans (marriages to admixed individuals, with some European ancestry, were still allowed though) (7). In early colonial times, mixed marriages were more acceptable than later on, and due to the population’s fast growth rate, early unions likely contributed exponentially more to the Afrikaner population. Elphick and Shell (3) distinguish two admixture patterns in Afrikaners based on historical records – in Cape Town and the surrounding area admixture was predominantly between European men and female slaves or former slaves, and in the outlying areas between European pastoralist frontier farmers (“trekboere”) and Khoe-San women.

Admixture with slaves (and former slaves) resulted from informal as well as formal associations (3). The church recorded many marriages between Europeans and manumitted slaves (7, 8). Husbands-to-be frequently had to buy their future slave wife’s freedom (7). It is unclear what the input of informal relationships into the Afrikaner gene pool was, as the outcome of these relationships and the population affiliation of the resulting offspring is not clear. Informal liaisons resulted, for example, from the slave lodge that served as a brothel for one hour a day for passing sailors and other European men (3, 22). This practice was so extensive that many children in the slave lodge clearly had European fathers (3/4 in 1671; 44/92 in 1685), prompting commissioner van Rheede to decree that these children should be manumitted and baptized (3). A census of the slave lodge in 1693 revealed 29/61 admixed school children and 23/38 admixed children under the age of three (3). Many women that married at the Cape during the early years used the toponym “van de Kaap” (meaning from the Cape) which may indicate a locally born slave. European men from slave-owning families also sometimes had a “voorkind” (meaning “before child”) with a slave in the household before they got married to a European woman (3). These children could also have been absorbed into the Afrikaner population (as opposed to becoming part of the Coloured population).

To understand the characteristics of the genetic contributions that slaves made it is necessary to know from where and when they came to Cape Town and see that in the light of European male partner choices. Shell (1994) (23) claimed that from 1658 to 1807, roughly a quarter of the slaves in the Cape colony came from Africa, Madagascar, South Asia and Southeast Asia each. Slave trade in the Cape was stopped in 1807 and slavery as such was stopped in 1834. Worden (24, 25) estimated that more slaves came from Asia, specifically South Asia, and fewer from Madagascar and Africa (supplementary text). Nevertheless, we do not expect an exact reflection of these ratios in Afrikaners. European men had a clear preference for Asian and locally born slaves over African and Madagascan women (3). Despite only two ships, containing west African slaves, that moored at the Cape in 1685 (26), we can expect the west African per capita contribution to exceed later arrivals because the fast population growth rate meant earlier contributions benefitted more from the exponential growth.

The “trekboere” were European farmers and their descendants followed a nomadic lifestyle in harsh conditions along the frontier. Informal unions with Khoe-San women were more frequent among the trekboere, but it is unclear if children from these relationships were absorbed into the Coloured and/or Afrikaner community (3, 4). Poor record keeping and a reduced presence of the church on the frontier meant that recorded information is incomplete for this section of the population. In the Cape, formal unions between European men and Khoe-San women were very unusual with only one known example (27).

By using church records, genealogists calculated the contribution of non-Europeans to be between 5.5% and 7.2% ((8-10) and supplementary text). These estimates may be biased because the registers; a) only reflect the Christian fraction of the population, b) were less complete at the frontier where admixture may have been more frequent, c) could be incorrectly pieced together from church records, and d) list people of unknown heritage, such as “van de Kaap”. In addition, records may be incorrect or unrecorded for children born out of wedlock. Populations that would have been excluded were a substantial Muslim community amongst manumitted slaves (3), a small Chinese population resulting from exiles and banned political prisoners (28, 29) and the indigenous Khoe-San who were not partial to the Christian religion (3). The presence of the Coloured population compounded these difficulties as genes may have exchanged between the Coloured and Afrikaner populations.

In order to clarify the patterns of ancestry and admixture fractions in current-day Afrikaners we compared genome-wide genotype data from 77 Afrikaners to comparative data of potential donor populations and tried to pinpoint the best possible sources of the admixture and the fractions of admixture from these groups.

## RESULTS

### Population structure and admixture

We generated filtered genotype data for 77 Afrikaner individuals (Method section) and merged the Afrikaner data with comparative data to create a dataset containing 1,747 individuals from 33 populations and 2,182,606 SNPs. We used this merged dataset to conduct population structure analysis and to infer population summary statistics.

In the population structure analysis (Figure 1) (30), Afrikaners cluster with non-Africans (K=2 to K=9) and specifically Europeans (K=3 to K=9) before receiving their own cluster at K=10. From K=7 onwards, northern and southern Europeans cluster separately, with Finnish forming one cluster (light blue) and southern Europeans (Tuscans and Iberians) the other cluster (light yellow). British (GBR) and Utah residents of northwestern European descent (CEU) appear midway between the Finnish and Southern European clusters. Afrikaners seem to contain slightly more northern European (blue) ancestry-component compared to CEU and GBR. The specific percentage of cluster assignments of Afrikaner individuals at K=6 and K=9, and the population averages assigned to each cluster, are given in Table S1 and S2.

**Figure 1:**
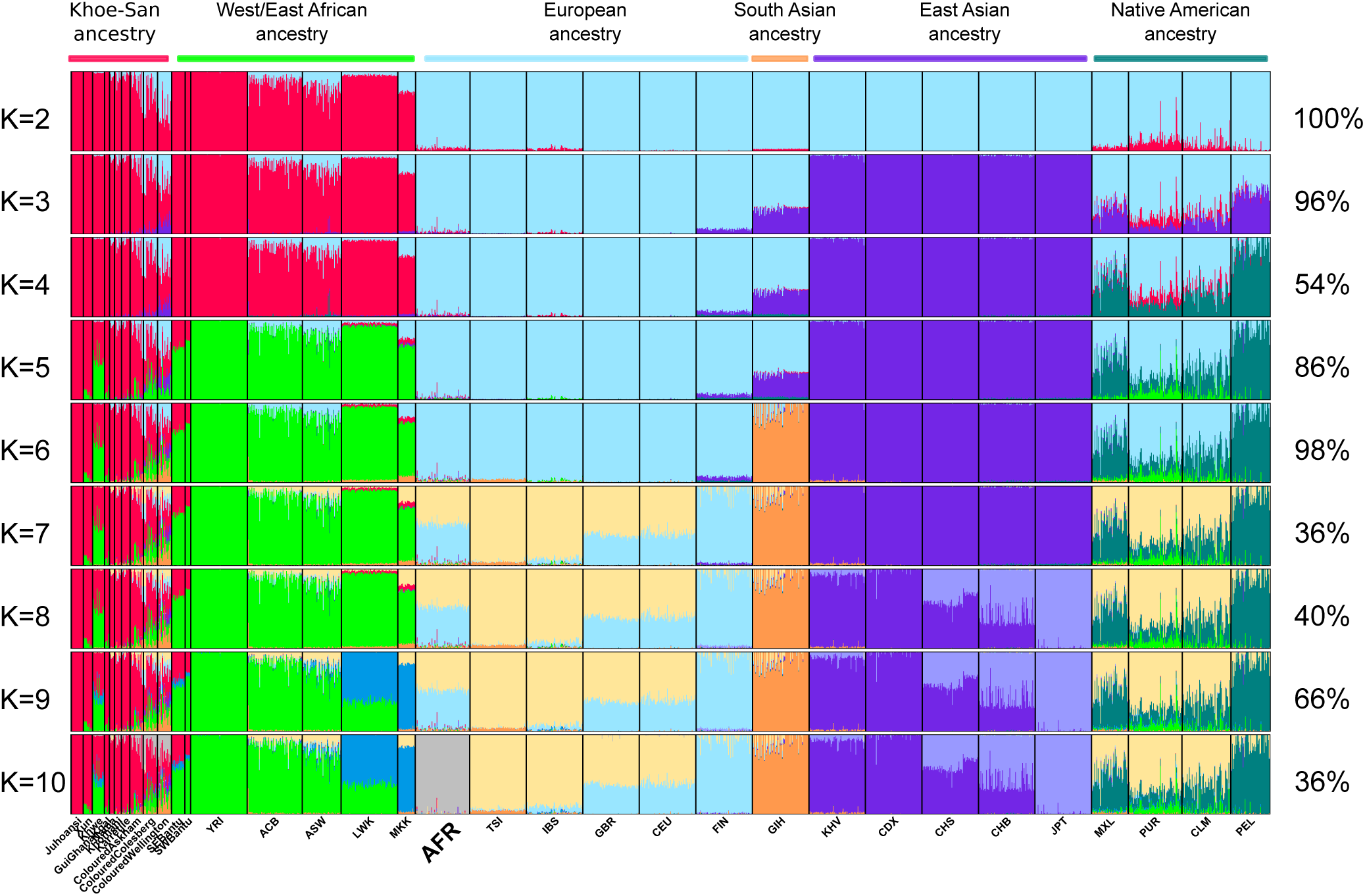
Admixture analysis of Afrikaners and comparative data. The number of allowed clusters are shown on the left, the statistical support on the right. Abbreviations: YRI-Yoruba from Nigeria, ACB and ASW-African American, LWK-Luhya from Kenya, MKK-Maasai from Kenya, AFR-Afrikaner from South Africa, TSI-Tuscan from Italy, IBS-Iberian from Spain, GBR-British from Great Britain, CEU-Northwest European ancestry from Utah, FIN-Finnish, KHV-Vietnamese, CDX-Dai from China, CHS-Han from southern China, CHB-Han from Beijing, JPT-Japanese, MXL-Mexican, PUR-Puerto Rican, CLM-Colombian, PEL-Peruvian.

Assuming six clusters (K=6), where the major geographical ancestries are discernible; i.e. aboriginal southern African (Khoe-San), West+East African, European, East Asian, South Asian, and Native American, the level of admixture from these ancestries can be distinguished in the Afrikaners (Table S1 and Figure 2). In addition to European ancestry (mean of 95.3% SD 3.8% - blue cluster) Afrikaners have noticeable levels of ancestry from South Asians (1.7% - orange cluster), Khoe-San (1.3% - red cluster), East Asians (0.9% - purple cluster), West/East Africans (0.8% - green cluster), and very low levels from Native Americans (0.1%). The small fraction from Native Americans likely stems from common ancestry between Native Americans and Europeans and from European admixture into Native Americans. The total amount of non-European ancestry, at the K=6 level, is 4.8% (SD 3.8%) of which 2.1% are African ancestry and 2.7% Asian/Native American ancestry. The individual with most non-European admixture had 24.9% non-European admixture and only a single Afrikaner individual (out of 77) had no evidence of non-European admixture (Table S1). Among the 77 Afrikaners investigated, 6.5% had above 10% non-European admixture, 27.3% between 5 and 10%, 59.7% between 1 and 5% and 6.5% below 1%.

**Figure 2:**
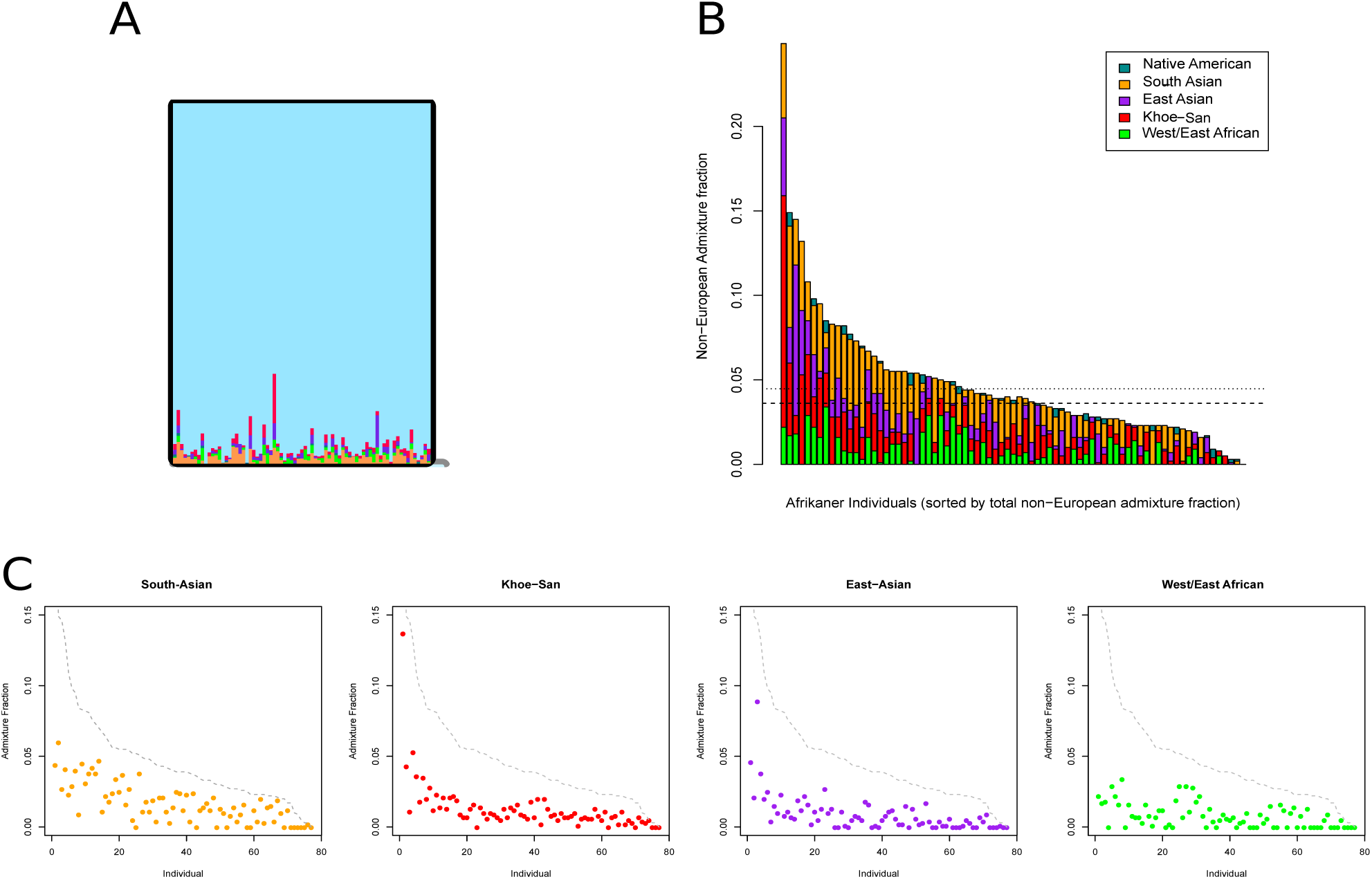
Admixture proportions of the Afrikaner at K=6. (A) Magnification of the Afrikaner population in the ADMIXTURE analyzes. (B) Non-European admixture fraction in the Afrikaner, sorted by total non-European admixture fraction. Dotted lines indicate the mean (top line) and median (bottom line) (C) Individual non-European admixture fractions sorted by total non-European admixture fraction (gray line).

The fractions of admixture from the different non-European groups in Afrikaners (at K=6) are generally correlated to each other (Figure S1), except for the West/East African admixture fractions.

At K=9 (before Afrikaners form their own cluster at K=10), additional inferences can be made regarding, Japanese vs. Chinese ancestry, East vs. West African Ancestry and northern vs. southern European ancestry (Table S2). Accordingly, it seems Afrikaners received East Asian ancestry from Chinese rather than Japanese individuals and slightly more West African ancestry than East African ancestry. Southern and northern European ancestry is almost equal in the Afrikaners but northern European ancestry is elevated compared to CEU and GBR.

Compared to the Afrikaners, the Coloured populations have more diverse origins. At K=6 the Cape Coloured population from Wellington (within the region of the original Cape colony) had the following ancestry fractions 30.1% Khoe-San, 24% European, 10.5% East Asian, 19.7% South Asian, 15.6% West/East African, and 0.2% Native American (Figure 1). The Coloured populations whom today are living further from the original Cape colony had different admixture patterns with less Asian and more Khoe-San contribution than the Cape Coloured: Colesberg Coloured (48.6% Khoe-San, 20% European, 2.9% East Asian, 6.7% South Asian, 21.6% West/East African, 0.2% Native American), Askham Coloured (76.9% Khoe-San, 11.1% European, 0.9% East Asian, 3.9% South Asian, 7.2% West/East African, 0% Native American).

In principle component analysis (PCA) (Figure 3 and S2) the first principal component (PC1) explains 3.6% of the variation in the dataset and distinguish Africans from non-Africans (right to left). PC2 explains 1.9% of the variation in the dataset and distinguishes Europeans from East Asians (top to bottom). The distribution of Afrikaners along PC1 and PC2 indicates both African and Asian admixture. Compared to northern Europeans (CEU and GBR) Afrikaners have more African and East Asian admixture. From the PCA it appears that most of the Afrikaner group have non-European ancestry at comparable levels to Iberians and Tuscans (IBS and TSI), however, certain Afrikaner individuals show greater levels of both African and Asian ancestry (Figure 3). Finnish populations show more East Asian but lower levels of African admixture than Afrikaners.

**Figure 3:**
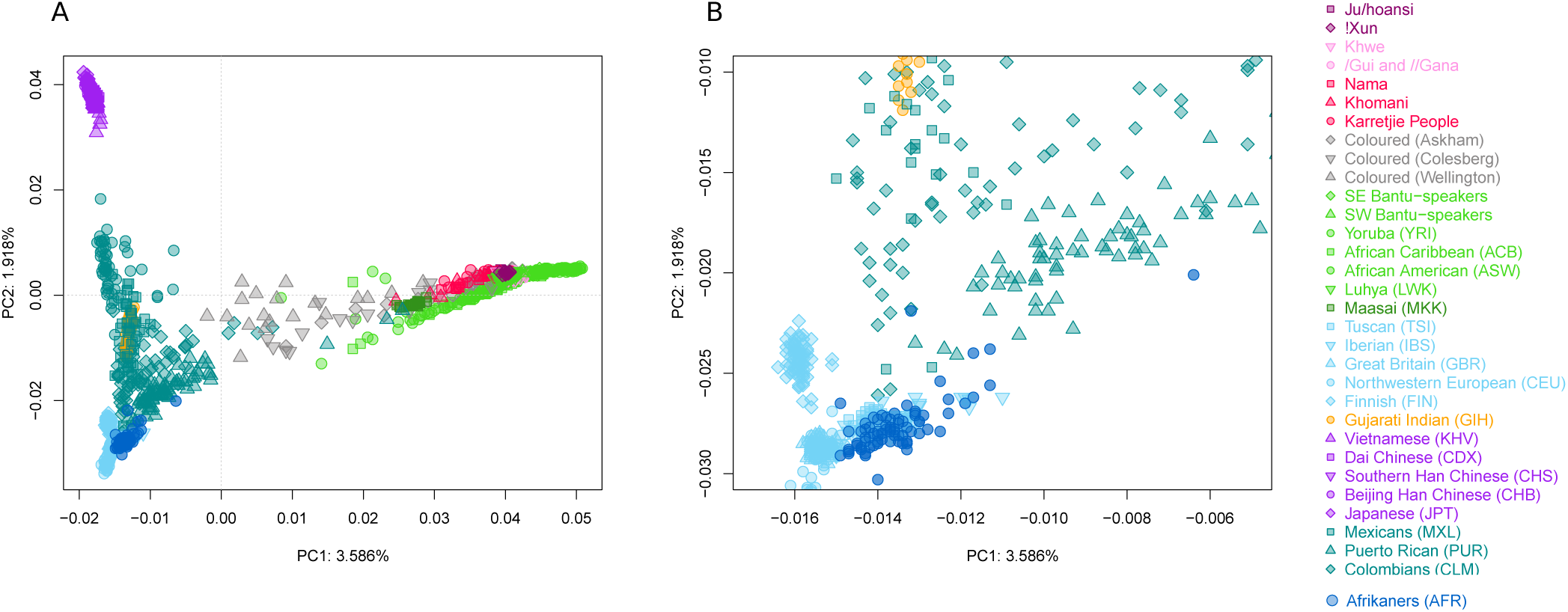
Principal component analyzes showing PC1 and PC2. (A) Full Figure, (B) shows a zoom-in on the Afrikaner (dark blue) population.

PC3 further defines variation within Africans and distinguishes between Khoe-San groups on the one extreme and groups with West African ancestry on the other. On this PC it appears that the African ancestry in Afrikaners has a greater affinity to Khoe-San variation (Figure S2). PC4 defines Native American ancestry and shows that the Afrikaner population is probably as similar to Native American populations as West European populations. PC5 delineates Asian ancestry and shows that the Afrikaner population may have genetic input from East Asian populations. PC6 distinguishes between West and East African ancestry and the Afrikaner population seems to be closer related to West African populations. PC7 differentiates between Japanese and Chinese ancestry, where we can confirm the affinity of the Afrikaner to the Chinese, as has been observed in the ADMIXTURE analysis. PC8 is associated with variation between different Khoe-San groups, where the Afrikaner cluster closer to the central and southern San than to the northern San. Further PC’s do not seem to capture group-wise information but continue to separate out individuals.

To look for the most likely sources of admixture in the Afrikaners within the comparative dataset we did formal tests of admixture (f_3_, (31)) between all pairs of comparative populations each time specifying (two) potential parental sources for the Afrikaner population. Results were sorted ascendingly according to Z scores (low Z scores indicate significant admixture) and Z-scores are visualized in a figure to aid interpretation (Figure S3), parental populations are colored according to regional population. It is clear that when only two populations are considered to be parental sources the most likely sources are always a European and either a Khoe-San or a West African population (combination of blue and red or blue and green labels in Figure S3). Subsequently the Coloured population can also be used to model a parental population in combination with Europeans (blue and grey labels). This method is however limited by the fact that only two parental sources can be tested at one time and might not be the best tests when multiple parental groups admixed into a population, as is the case for the Afrikaner population.

We also used projected PCA analysis (32) by constructing principal components (PCs) based on certain specified populations and then projecting Afrikaners on existing PC’s to further refine specific ancestry components within bigger ancestral groups (Figures S4 and S5). To distinguish the specific Khoe-San group who contributed to Afrikaner variation, we constructed a PCA based on the variation of Karretjie People (Southern San), Ju|’hoansi (Northern San) and CEU (northwestern Europeans) and projected Afrikaners on the PCA. The projection (Figure S4A-B), clearly shows that Afrikaners have more Khoe-San admixture than CEU (see shift in the first PC) and furthermore, that the admixture was from a Khoe-San population is more similar to Karretjie People (southern San) than Ju|’hoansi (northern San) as observed in the skew towards Karretjie. This trend towards southern San rather than northern San is even made clearer when YRI is used in place of CEU in the projection (Figure S4C-D). This shift of AFR in PC1 is not only due to drift and inherent population differences between populations of European origin, because when GBR is added to the projection, GBR and CEU overlap with each other, with AFR still clearly shifted towards Khoe-San (Figure S4E-F). Other southern Khoe-San groups (≠Khomani and Nama) show similar behaviors in terms of being favored compared to northern San (Figure S5).

Alleles that are shared privately between combinations of the Afrikaner population with one comparative population (Figure S6) show that the Afrikaner share the most private alleles with the CEU population, which makes them a better parental source than the other European populations. The ≠Khomani share the most private alleles with the Afrikaner out of all Khoe-San populations, which is consistent with the previous observations. The most shared private alleles of the Afrikaner with Asian populations can be found in the GIH (Gujarati Indian) followed by the CHB (Chinese) and JPT (Japanese). The Afrikaner share more private alleles with the Niger-Congo speaking Yoruba (YRI) from Nigeria than with the two eastern Bantu-speaking groups, Luhya (LWK) from Kenya and southeastern Bantu-speakers from South Africa (Figure S7). This supports an admixture from West-African slaves rather than from southern African Bantu-speakers into the Afrikaner population.

For finer scale resolution of the European and the Asian components in the Afrikaners, the dataset analyzed above was furthermore merged, respectively, with the POPRES dataset (33, 34) and various datasets containing populations from East, South and Southeast Asia. These additional European and Asian comparative datasets had much lower SNP densities but they contained many more comparative populations and were used for fine scale resolution of European and Asian components in Afrikaners.

The Afrikaner individuals were projected on a PCA, constructed based on European variation present in the POPRES dataset (Figure S8). Afrikaner individuals seem to group within western European variation. They are grouped in-between the French and German clusters on the PC plot. When principal components were summarized by population averages and standard deviations – they seem to be grouping closest to Swiss German, Swiss French and Belgian populations. This positioning could also be explained as an intermediate position between German, Dutch and French variation that link to each other in a clinal pattern (33, 34).

When analyzing Afrikaners with a dataset enriched for Asian populations it appears that the largest contributing Asian component is from India (Figure S9). The orange component in Figure S9 is the most prominent admixed component from Asian groups and this component is specifically associated with Indo-European speaking Indian groups, i.e. Khatri, Gujarati Brahmin, West Bengal Brahmin, Maratha; and Dravidian speaking Iyer (35).

### Genome local ancestry and selection

To see if non-European ancestries were enriched in any specific part of the genome, we also inferred local genomic ancestries in Afrikaner individuals (Figure S10B). Specific enrichment of admixed ancestries in a specific region of the genome could be indicative of a possible adaptive introgression event. There are two regions of interest, one on chromosome 2 and another on chromosome 21, both enriched for Khoe-San ancestry. For the region on chromosome 21 between positions 10,699,687-14,537,654 Afrikaners are 100% assigned to Khoe-San ancestry (Fig S10B). The three genes associated with the region, *TPTE, BAGE2* and *ANKRD30BP2*, all have exclusive expression in the testis. The *TPTE* gene encodes a tyrosine phosphatase which may play a role in the signal transduction pathways of the endocrine or spermatogenic function of the testis (UCSC genome browser). The *BAGE2* gene is from the B melanoma antigen family and is not expressed in normal tissues except the testis. It is however also expressed in 22% of melanomas and have been associated with Melanoma skin cancer (UCSC genome browser). *ANKRD30BP2* is a non-coding RNA pseudogene with exclusive expression in the testes (UCSC genome browser). This region on chromosome 21 span the centromere of the chromosome, which could influence SNP assignment, and results should be interpreted with caution (FigS10B). The region on chromosome 2 shows around 10% Khoe-San ancestry and is located between positions 2,974,989 – 3,058,838 in the genome. This region is located close to the telomere of chromosome 2, which might influence results. There is one annotated human cDNA that spans this region, namely *AK095310*. The gene function is not known but the highest expression is in the brain, with some expression in the pituitary glands (UCSC genome browser).

We also analyzed the Afrikaner data for genome-wide signals of selection by scanning for regions with extended haplotype homozygosity compared to other haplotypes within the same population (iHS scans) and compared to haplotypes in a comparative population (XP-EHH scans). For XP-EHH scans, Afrikaners were compared to the CEU population. Figure S11 shows the genome-wide Manhattan plot of selection scan results for iHS and XP-EHH. Several peaks that might indicate signals of selection in the Afrikaner group were observed. The top 5 peaks in each scan are listed in Table S3 together with; position information, underlying/nearby genes and potential function of underlying/nearby genes. From the top 5 iHS peaks only one had a gene directly associated with the peak, the gene *FGF2* is a fibroblast growth factor with a variety of functions and was previously associated with cholesterol levels. The XP-EHH results were clearer and three out of the five top peaks were directly associated with genes; *CCBE1* – a gene previously associated with lymphatic disease, *ACTG2* – an enteric smooth muscle actin gene previously associated with intestinal diseases and *SUCLG2* – encoding a succinate-CoA ligase, previously associated with glucose and fat metabolism. Interestingly the Afrikaner group does not show the strong adaptation signals at the lactase persistence region on chromosome 2 and MHC region on chromosome 6, which is strong and well-known for the CEU group. Although the CEU group have significantly more of the lactase persistence associated allele (rs4988235-T) (71% in CEU vs. 52% in the AFR, *p*= 0.000434), their predicted lactase persistence phenotypic status, based on homozygote and heterozygote counts combined, is not significantly different (91% in CEU vs. 83% in AFR, *p*= 0.126514).

### Dating of admixture

We constructed linkage disequilibrium (LD) decay curves, to date the various admixture events in the Afrikaners. Clear LD decay curves are visible, indicating admixture, when CEU and an admixing group are used as parental populations and Afrikaners as admixed population (Table S4, Figure S12). A date of 9-10 generations ago was obtained with African populations as the second parental population and around 15-16 generations ago with Asian populations as second parental population.

### Estimation of bottleneck effects

Runs of homozygosity were calculated for each individual of the dataset. Depending on their length, runs of homozygosity are informative of historic population size or recent inbreeding in populations (36). While we see striking differences between continental groups (Figure 4), there is no strong difference between the Afrikaner and other European populations, except for the Finish population that appears to have had a smaller historic effective population size (Figure 4). The average inbreeding coefficients were also calculated for all populations (Table S5) and Afrikaners had a lower average inbreeding coefficient than other European populations.

**Figure 4:**
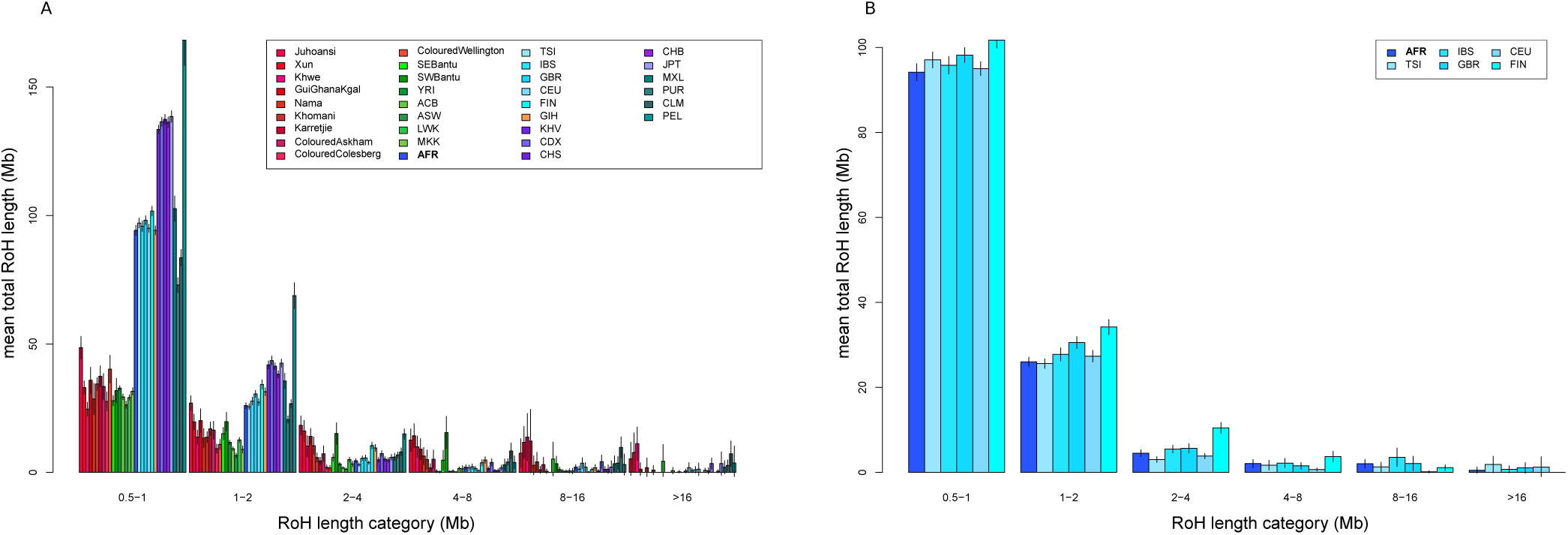
(A) Runs of Homozygosity. Populations are colored according to regional group. (B) Magnification of runs of homozygosity for European populations and the Afrikaner.

## DISCUSSION

Genealogical records suggest that Afrikaners have their main ancestry components from Europeans (Dutch, German and French) and estimate the non-European contributions to the Afrikaner to be between 5.5% and 7.2% ((8, 9) and supplementary text). Our genetic study that included 77 Afrikaners, inferred a slightly lower non-European contribution than predicted by genealogical studies. From population structure analyzes we saw that Afrikaners have their main ancestry component (95.3%) from European populations. The European component is a more northwestern (than southern or eastern) European component (Figure 1 and S8), which is in agreement with genealogical records of most ancestry coming from Dutch and German (61-71%), intermediate from French (13-26%), with much smaller fractions from other European groups ((8) and supplementary text). Of note, Afrikaners group separately from populations from the UK (Figure S8) despite the fact that the Cape was a British colony from 1806 onwards. This confirms the relatively small contributions from British people to the Afrikaner population as predicted by genealogical records (8).

The non-European fraction in Afrikaners was estimated to 4.7% on average (Table S1). More of the non-European admixture fraction appeared to have come from people who were brought to the Cape as slaves (3.4%) during colonial times than from local Khoe-San people (1.3%). Indeed historical records of the early Cape Colony record more instances of unions between European men and slaves or former slaves than to local Khoe-San women (3). Only one example of a Khoe-San - European union in the Cape colony is known. A local Khoekhoe woman from the Goringhaicona group, Eva (or Krotoa) van de Kaap was an interpreter and ambassador between the colonists and Khoekhoe people and married Pieter Van Meerhof in 1664 (27, 37). Since unions between Khoe-San women and the frontier farmers were thought to be more frequent, it may account for the 1.3% observed admixture in the Afrikaner population. The 1.3% observed Khoe-San ancestry calculates to 26.6 Khoe-San women out of 2048 ancestors 11 generations ago. However, we know that one Afrikaner had for example only 299 ancestors in colonial times (10) because many Afrikaner ancestors enter pedigrees multiple times (8, 10, 38). These 26.6 Khoe-San women that contributed to the average Afrikaner should thus not be seen as 26 separate women (i.e. the same woman could have contributed many times). The Khoe-San admixture component is the most ubiquitous non-European admixture component and only 6 out of the 77 Afrikaners had no Khoe-San ancestry (Table S1). The Khoe-San ancestry in Afrikaners appears to have come from Southern Khoe-San groups and not Northern Khoe-San (Figure S5 and S6). This is not surprising since in pre-historic and historic times, the Southern Khoe-San inhabited current-day South Africa while the Northern San lived in northern Namibia and southern Angola (12, 16, 39-41).

South and East Asians contribute cumulatively to an average of 2.6% of the Afrikaner ancestry (53.2 out of 2048 ancestors 11 generations ago). Elpick and Shell (1989) (3) noted that European men more often mixed with Asian and locally born slaves than African and Madagascan women. Although many other additional factors might have played a role in the resultant current-day Afrikaner admixture fractions, the genetic admixture fractions of South and East Asians was higher in current-day Afrikaners than Khoe-San fractions (1.3%) and West/East African fractions (0.8%), and slightly higher than the combined African fraction (2.1%). South Asian contributions outweigh East Asian contributions (p-value of <0.00001, paired Wilcox test) (Table S2). The South Asian contribution seems to have come predominantly from Indian populations (Figure S9).

West/East Africans contributed an average of 0.8% of the Afrikaner ancestry (Table S1) (16.3 out of 2048 ancestors 11 generations ago). Shell (23) estimated that about 63,000 slaves arrived in the Cape colony between 1658-1807 and a quarter came from West/East coastal Africa (26.4%, east coast and only 2.5% from West Africa). Only two ships brought West African slaves to the Cape in 1685 (26). When one takes into account that only 2.5% of African slaves came from West Africa, it is fascinating that just over half of this signal is from West Africans rather than East Africans (Table S2). This discrepancy could possibly be explained by West Africans arriving four generations earlier than East Africans (see Supplementary text). More frequent admixture during early years, and fast population growth could have caused the genetic footprint of West Africans to exceed that of East Africans. While admixture fractions between East Asians, South Asians and Khoe-San correlate well with each other in Afrikaner individuals (Figure S1), West/East African fractions do not correlate significantly with South Asian and East Asian fractions and a high number of Afrikaners had no West/East African admixture (26/77). These patterns could possibly be explained by the fact that, there were relatively few West African slaves at the Cape, that the arrival of West African slaves was contained in a very limited time period, and that East African slaves arrived later in time. The West African fraction in the Afrikaners is presumed to come mostly from these West African slaves and not from southern African Bantu-speakers. This is confirmed by shared allele analysis (Figure S7) where Afrikaners share more alleles with the West African Yoruba from Nigeria than with local South African Bantu-speakers (southeast Bantu-speakers). Although current-day South African Bantu-speakers trace the majority of their ancestry (80%) to West Africa (12, 16, 40, 42) (Figure 1), there were no Bantu-speakers present in this part of southern Africa during colonial times (22). Bantu-speakers originated from West Africa and started to expand to the rest of sub-Saharan Africa around 4,000 years ago, reaching the northern parts of current-day South Africa around 1,800 years ago. During the 1600s the edge of the Bantu-expansion (specifically Xhosa speakers) reached as far south as the Fish River in the Eastern Cape Province of current day South Africa (approximately 1000 km from Cape Town). Therefore, Bantu-speakers were only encountered at a much later stage during the frontier wars in 1779-1851 by expanding Afrikaner frontier farmers (22). Although initial interactions between the frontier farmers and Bantu-speaking Xhosa populations were peaceful with trade between groups, relations quickly turned aggressive and conflicts erupted. It is unlikely that much gene flow occurred between South African Bantu-speakers and Afrikaners, since during this relatively late time period, the Afrikaner population identity had already culminated, which resulted in social hinders to relationships between groups.

Genome local ancestry analysis revealed two possible adaptive introgression signals in the Afrikaners, on chromosome 2 and 21. In both cases Khoe-San ancestry was enriched for these genomic regions. The chromosome 21 region was 100% assigned to Khoe-San ancestry and the three genes in this region are exclusively expressed in the testes and might indicate an adaptation linked to reproduction or male fertility in the Afrikaners. Since the implicated regions are close to centromeres or telomeres results should be interpreted with caution. Scans for selection in the Afrikaners implicated several genes associated with diet, i.e. intestinal function, lipid and glucose metabolism, possibly indicating adaptation to modified or novel food sources.

The estimated times of admixture differs between African and Asian populations (Table S4) and was estimated for around 9-10 generations for West Africans and Khoe-San and around 15-16 generations for Asians. In the Afrikaner population the average generation time for men was 32.92 years whereas for women it was 27.36 (43), suggesting that the Afrikaner population is on average between 11 and 13 generations old. Since many admixing events occurred during early colonial times, genetic dating of admixture times is expected to fall in this range as well. The admixture times for Asian populations appeared older though. Some possible explanations for this could be that some of the slaves may already have been admixed when they arrived so that a generation or two could be added; previous admixture into Europeans might play a role; there could have been more recent admixture events with Africans in Afrikaners that could contribute to more recent dates; the differences in effective population sizes between Africans and non-Africans might influence estimates.

It is interesting to observe that Afrikaners do not present a signal of a population bottleneck compared to European groups even though they had a very small founding population (Figure 4 and Table S5). This could be explained by the fact that even though the initial founding population of the Afrikaners was small, they were from diverse origins and many of the initial unions resulted in admixed children who were incorporated in the resultant population. For example, one Afrikaner individual (JMG - (10)) had 125 common ancestors, but these were so distant from each other that his inbreeding coefficient is only 0.0019. Until recently, most humans were sedentary and populations were small so that inbreeding due to distant relations was not unusual. However, a number of founder effects for specific diseases have been identified in Afrikaners ((44, 45) and supplementary text). These founder effects however need not imply inbreeding but rather suggest a sampling effect, i.e. some disease alleles were present in original founders and were amplified through exponential population growth.

## CONCLUSION

Although Afrikaners have the majority of their ancestry from northwestern Europe, non-European admixture signals are ubiquitous in the Afrikaner population. Interesting patterns and similarities could be seen between genealogical predictions and genetic inferences. The non-European admixture fractions resulting from people that were brought to southern Africa as slaves and local Khoe-San populations might have been one of the factors that helped to counteract the adverse effect of a small founding population size and inbreeding, since Afrikaners today have comparable inbreeding levels to current-day European populations.

## MATERIALS AND METHODS

### Sample collection and genotyping

Ethical clearance for this study was obtained from the ethics committee from the Natural and Agricultural Science of the University of Pretoria EC11912-065. All subjects were given a study information sheet and gave their written informed consent for the collection of samples and subsequent analysis.

The 77 individuals included in this study form part of parallel studies on non-paternity (43, 46) and on the mitochondrial DNA heritage of self-identified Afrikaners. Fifty three samples came from 17 groups of men bearing the same surnames (an average of 3.12 individuals per family with the same surname). Males with the same surname are separated by an average of 15.8 generations along their patrilines. Twenty four samples are from unrelated patrilines, either having unique surnames or stemming from different founders.

Samples were collected with Oragene® DNA Saliva collection kits (DNA Genotek, Kanata, Canada) and whole genome DNA was extracted according to the manufacturer’s instructions. Final concentrations were adjusted to 50 ng/µl. Genotyping was performed by the SNP&SEQ Technology Platform in Uppsala, Sweden (www.genotyping.se) using the Human Omni 5M SNP-array. Results were analyzed using the software GenomeStudio 2011.1, and the data were exported to Plink format and aligned to hg19.

### Genotype filtering and merging with Comparative data

SNP data quality filtering and merging to comparative data was done with PLINK v1.90b3 (47). A 10% genotype missingness threshold was applied and the HWE rejection confidence level was set to 0.001. SNPs with a chromosome position of 0, indels, duplicate-, mitochondrial-and sex chromosome SNPs were removed. All individuals passed a missingness threshold of 15% and a pairwise IBS threshold of 0.25 (for identification of potential relatives).

The resultant dataset of 4,154,029 SNPs and 77 individuals were phased using fastPHASE (48), with 25 haplotype clusters, 25 runs of the EM-algorithm, and 10% assumed missingness. Subsequently the data was merged with data from (16), containing 2,286,795 quality filtered autosomal SNPs typed in 117 southern African Khoe-San and Bantu-speakers. Before merging the datasets, AT and CG SNPs were removed from the datasets. During the merge the strands of mismatching SNPs were flipped once and remaining mismatches were removed, only the overlapping positions between the datasets were kept.

To get a more extensive set of African and non-African comparative data, we furthermore downloaded SNP data from the 1000 Genomes Project website, at ftp.1000genomes.ebi.ac.uk/vol1/ftp/technical/working/20120131_omni_genotypes_and_intensiti es (49). The 1000 genomes genotype data were quality filtered using the same thresholds as used in our datasets (described above). The following populations were included from the 1000 genomes dataset: YRI and LWK (Yoruba and Luhya - West African ancestry), MKK (Maasai - East African), ACB and ASW (African-Americans in the Caribbeans and southwest USA), TSI, IBS, CEU, GBR, and FIN (Tuscans, Iberians, northwest European ancestry, British, Finnish – European), JPT, GIH, CHB, CHS, CDX, KHV (Japanese, Indian ancestry, Han Chinese Bejing, Han Chinese South, Dai Chinese, Vietnamese – Asian), PEL, PUR, MXL, CLM (Peruvians, Puerto Ricans, Mexican ancestry, Colombians – Native American (admixed)). All populations were randomly down-sampled to 80 individuals. This merged dataset included a total of 2,182,606 high quality SNPs in 1,747 individuals from 33 populations.

For finer scale resolution of the European component and the Asian component in Afrikaners, this dataset was furthermore merged with 1) the POPRES dataset (33, 34) and 2) various datasets containing populations from east, south and southeast Asia (35, 50-56). These European and Asian comparative datasets were quality filtered and phased with the same thresholds and parameters as used in the previous datasets. Although these datasets had much lower SNP densities (149,365 SNPs for the European and 313,790 SNPs for the Asian dataset) they contained many more comparative populations (37 European comparative populations for the European dataset and 63 Asian comparative populations for the Asian dataset).

### Population structure analysis

Population genetic analysis was conducted for the main merged dataset, containing 1747 individuals from 33 populations and 2,182,606 SNPs. We inferred admixture fractions (30) in order to investigate genomic relationships among individuals based on the SNP genotypes. Default settings and a random seed were used. Between 2 and 10 clusters (K) were tested. A total of 100 iterations of ADMIXTURE were run for each value of K and the iterations were analyzed using CLUMPP (57) for each K to identify common modes among replicates. Pairs of replicates yielding a symmetric coefficient G’>=0.9 were considered to belong to common modes. The most frequent common modes were selected and visualized with DISTRUCT (58). For the Asian extended dataset similar settings were used as described above, however clustering was done for K=2 to K=15 due to the higher number of populations in the dataset.

PCA was performed with EIGENSOFT (32, 59) with the following parameters: r2 threshold of 0.1, population size limit of 80, and 10 iterations of outlier removal. Projected PCA analysis was done using EIGENSOFT by constructing Principal Components (PCs) based on certain specified populations and then projecting Afrikaners on existing PC’s.

Formal f3 tests of admixture (31) were done between all pairs of comparative populations specifying (two) potential parental sources of the Afrikaner population. We estimated the admixture time based on linkage disequilibrium (LD) decay due to admixture (ROLLOFF - (31)), default parameters were used. The standard error was estimated with a jackknife procedure. Generations were converted to years using 29 years per generation. Shared private alleles were inferred using ADZE (60) for all pairwise population combinations of populations with at least 15 individuals. Runs of homozygosity were calculated using PLINK (47) using the following parameters (--homozyg --homozyg-window-kb 5000 --homozyg-window-het 1 --homozyg-window-threshold 0.05 --homozyg-kb 500 --homozyg-snp 25 --homozyg-density 50 --homozyg-gap 100). Inbreeding coefficients were calculated using PLINK (--het).

### Local ancestry analysis

We inferred genome local ancestry for the Afrikaner individuals using RFMix version 1.5.4 (61). The following populations were used as putative sources: CEU, CDX, YRI and Khoe-San groups (combined !Xun, ≠Khomani, Karretjie and Ju|huansi). RFMix was run with the following settings: RFMix_v1.5.4/RunRFMix.py --forward-backward -e 2 infilename. Other settings were left as default. The results were visualized using the ancestry pipeline by Alicia Martin available at https://github.com/armartin/ancestry_pipeline/blob/master/plot_karyogram.py and a Python plotting script making use of the plotting library Bokeh. The first 2 Mbp from each end of the chromosomes were filtered to avoid telomere regions.

To scan for signals of genome-wide selection in the Afrikaner group, integrated haplotype scores (iHS) and the cross population extended haplotype homozygosity (XP-EHH) were analyzed using R package REHH (62). The ancestral state was identified by its presence in the chimpanzee, gorilla, orangutan and human genomes (downloaded from UCSC). Based on this requirement, we performed selection analyzes on 1,759,008 SNPs. iHS and XP-EHH were calculated with maximum distance between two SNPs of 200,000 bp. For the XP-EHH we compared the Afrikaners (AFR) haplotype homozygosity with Northwest European ancestry individuals (CEU).

### Data availability

The anonymized genome-wide SNP data of 75 of the 77 Afrikaner individuals who consented to have their data shared electronically, will be made available for research use through the ArrayExpress database (https://www.ebi.ac.uk/arrayexpress), access number XX.

## ACKNOWLEDGEMENTS

We thank Karen Harris, Michele Ramsay, Cesar Fortes-Lima and Maximillian Larena for their useful comments and Nigel Worden for invaluable information. We thank all the participants in the study. The POPRES data were obtained from dbGaP (accession no. phs000145.v1.p1). This work is based upon research supported by the National Research Foundation of South Africa (grant 77256 to JMG) the Genomics Research Institute of the University of Pretoria (to JMG) and by the Swedish Research Council (no. 621-2014-5211 to CS and 642-2013-8019 to MJ) and Knut and Alice Wallenberg foundation (to MJ). JCE was supported by an NRF scarce-skills PhD scholarship, a University of Pretoria study abroad bursary and bursary allocations from JMG.

