## Supplement for "Patterns of African and Asian admixture in the Afrikaner population of South Africa"

Running Title: Admixture into the Afrikaner population of South Africa

* Equal contribution
[+] Equal contribution

[#] Correspondence: Carina Schlebusch


[#] Correspondence: Jaco Greeff


Keywords: Afrikaner, South Africa, admixture, slave trade, colonial times

### SUPPLEMENTARY FIGURES AND TABLES

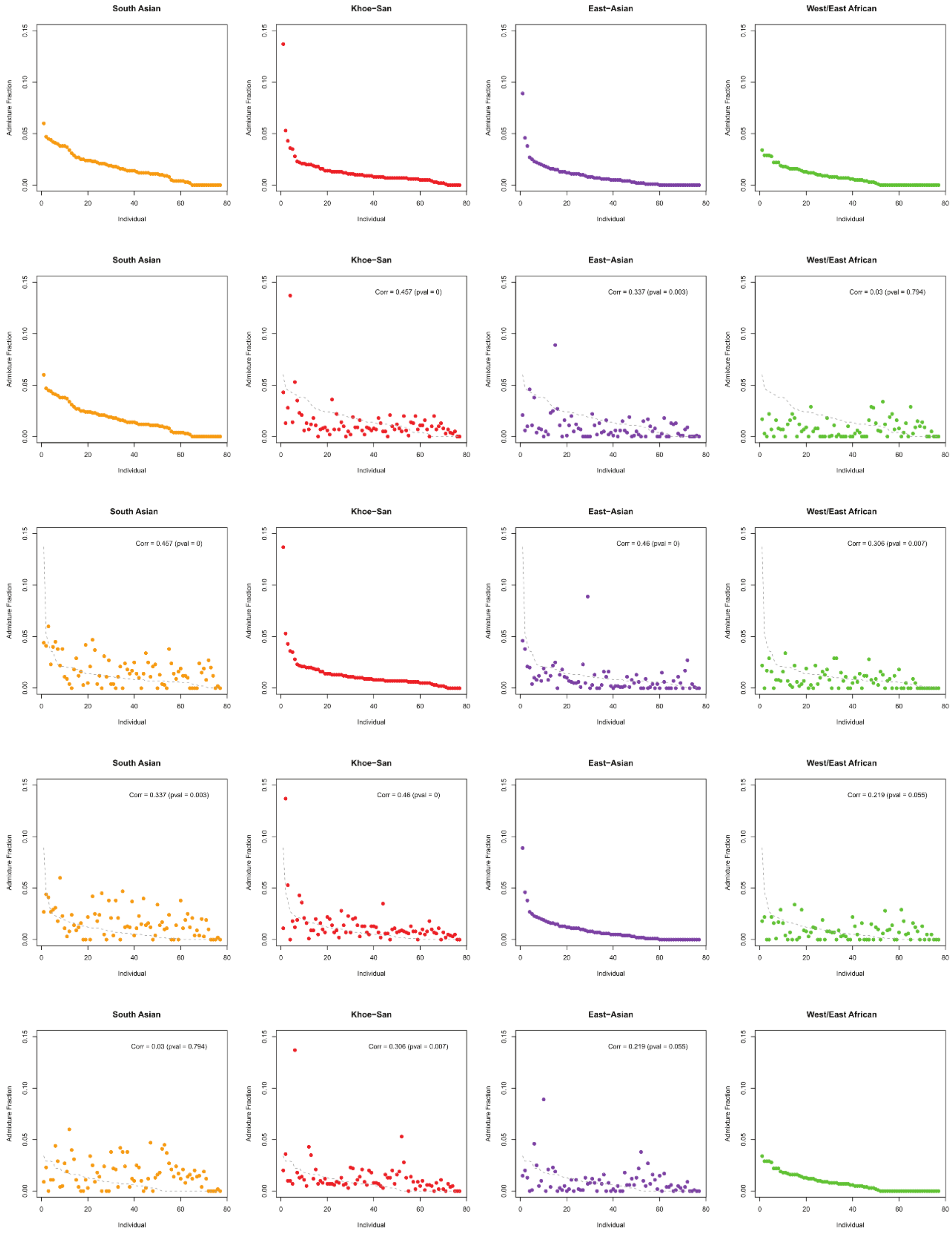


*Figure S1: Non-European admixture fractions (of K=6) sorted by ancestry fraction. The first row shows all admixture fractions sorted by their own ancestry, the next row is sorted according to South Asian ancestry, then Khoe-San ancestry, East Asian ancestry, and finally West African ancestry. Ancestry correlations and significance of correlation are indicated on graphs.*

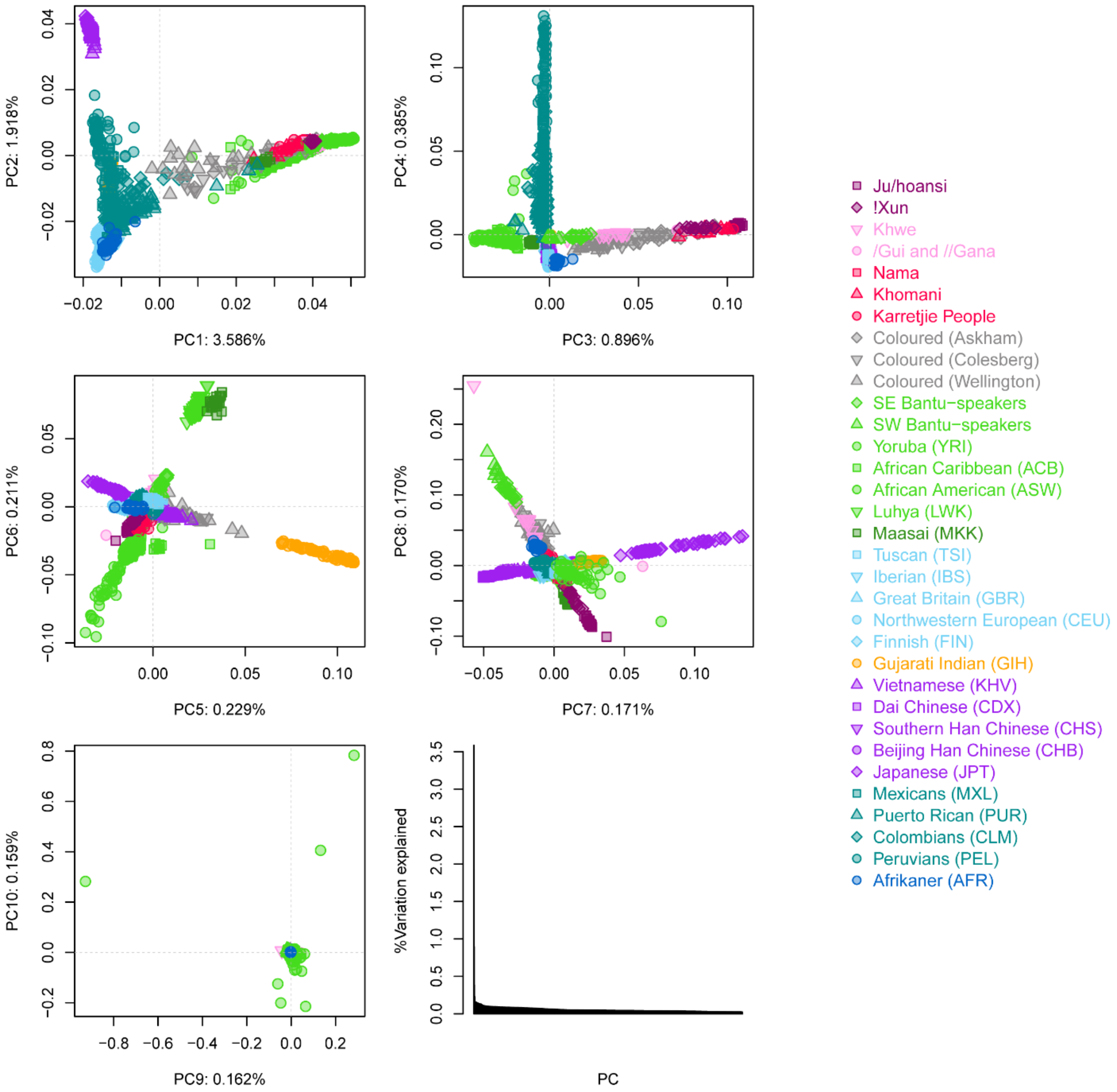


*Figure S2: Principal component analysis for PC1-PC10 and the variation explained by PCs. Populations are colored according to regional grouping.*

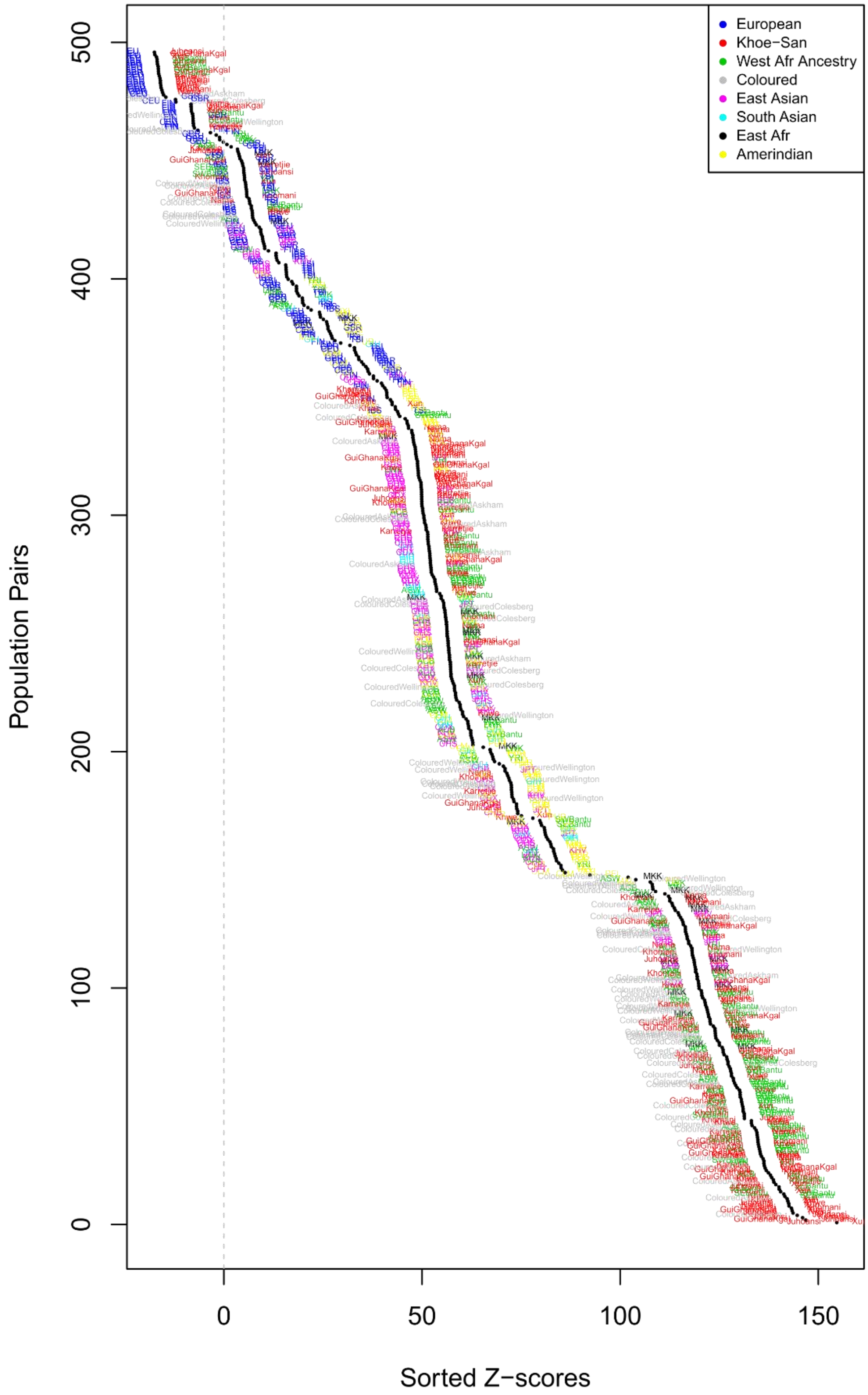


*Figure S3: Results from $f_3$-test. Populations are colored according to regional affiliation.*

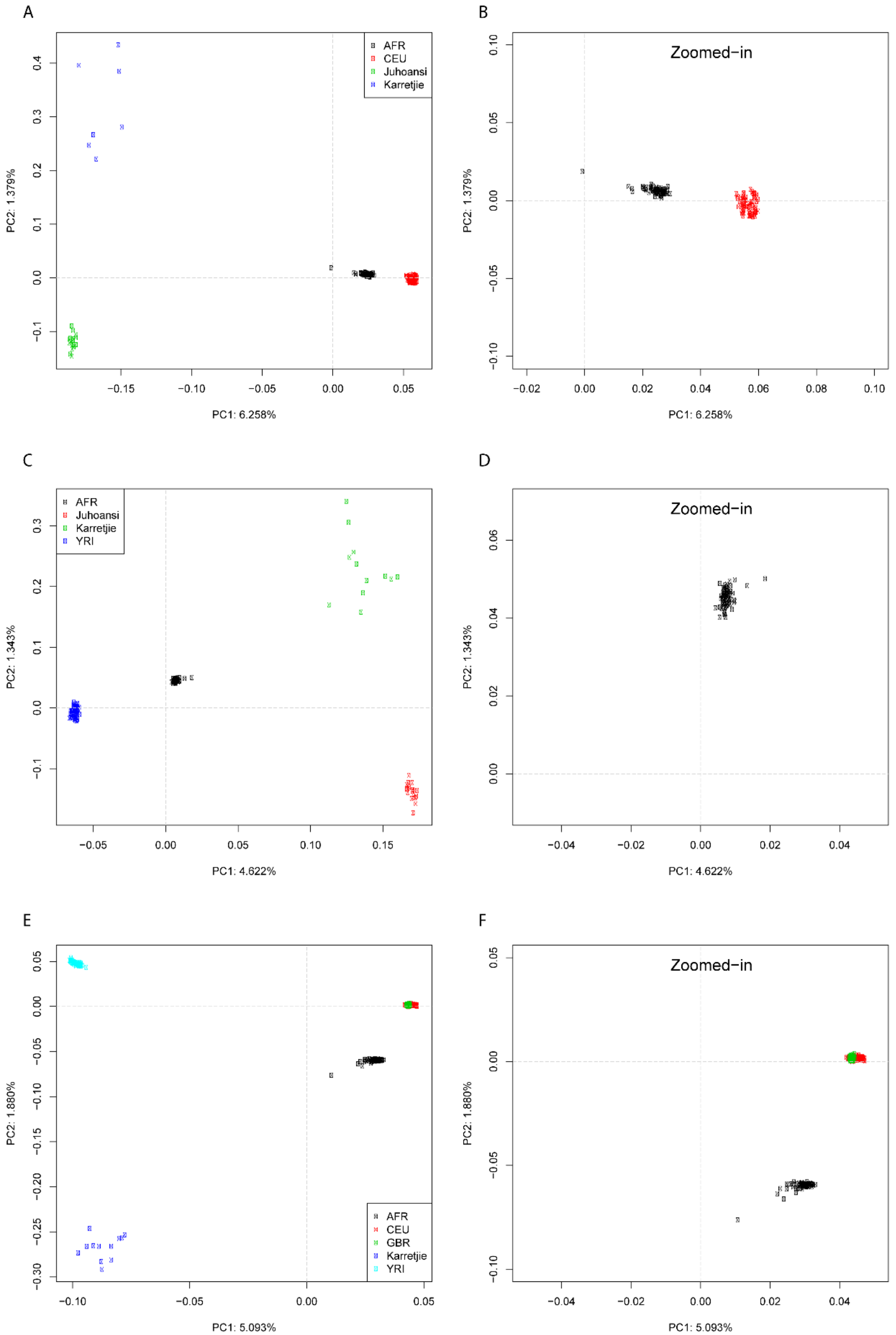


*Figure S4: Projected PCA. (A, B) Afrikaner population was projected on a PCA of CEU, Juhoansi, and Karretjie. (C, D) Afrikaner population was projected on a PCA of YRI, Juhoansi, and Karretjie. (E, F) Afrikaner population was projected on a PCA of CEU, GBR, YRI, and Karretjie.*

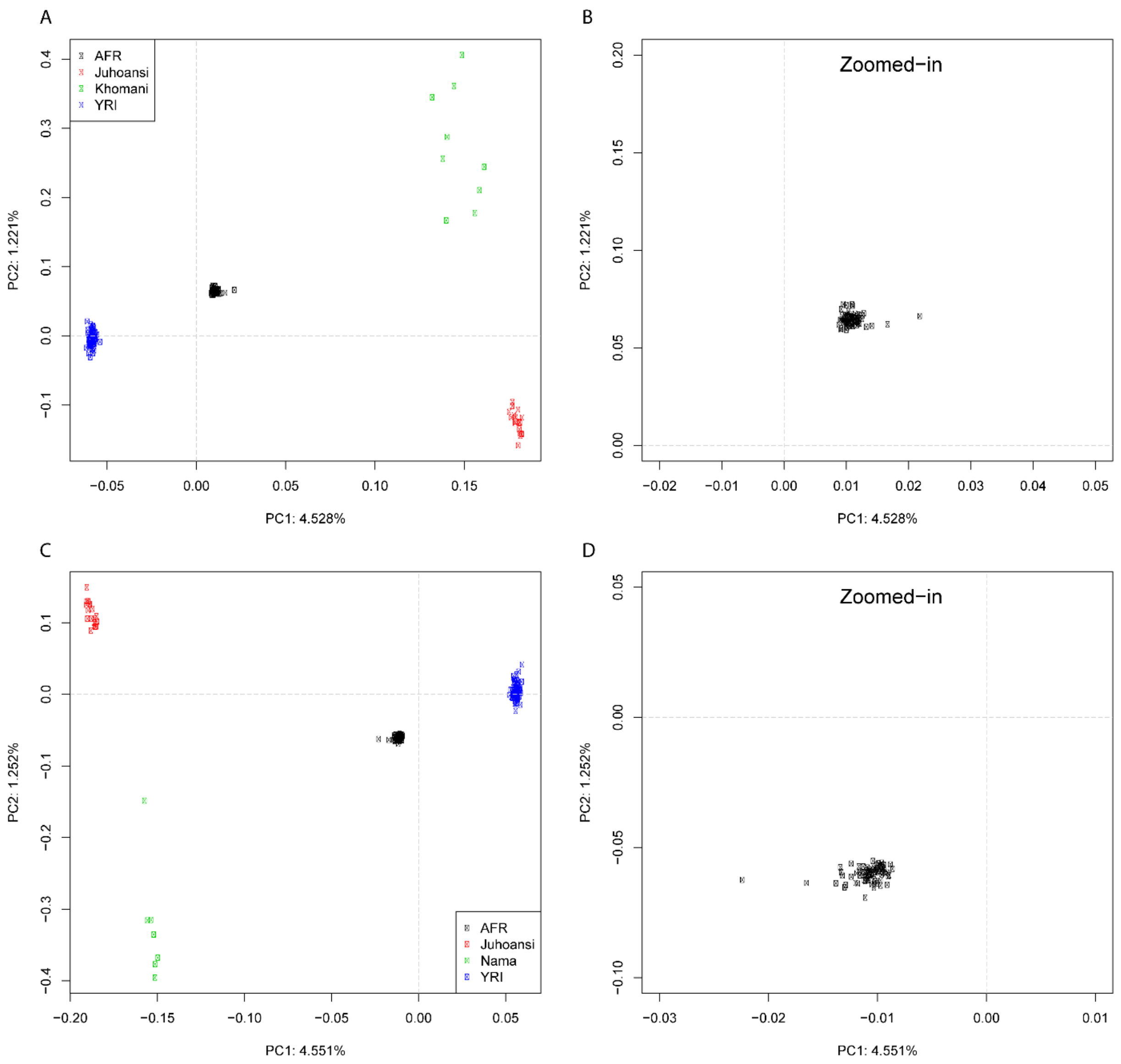


*Figure S5: Projected PCA. (A, B) Afrikaner population was projected on a PCA of YRI, Juhoansi and Khomani. (C, D) Afrikaner population was projected on a PCA of YRI, Juhoansi and Nama.*

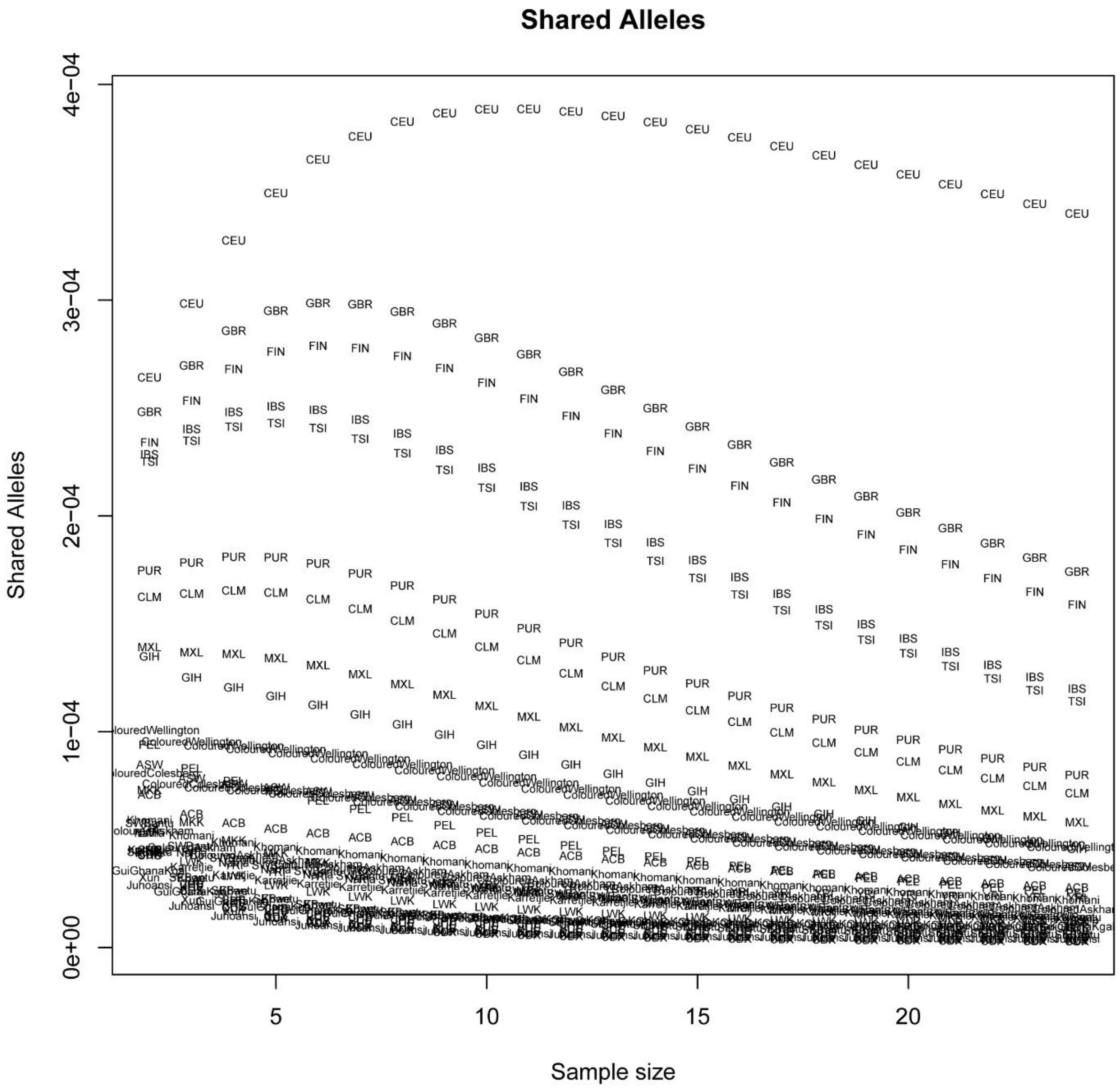


*Figure S6: Fraction of shared private alleles between the Afrikaner population and a comparative population.*

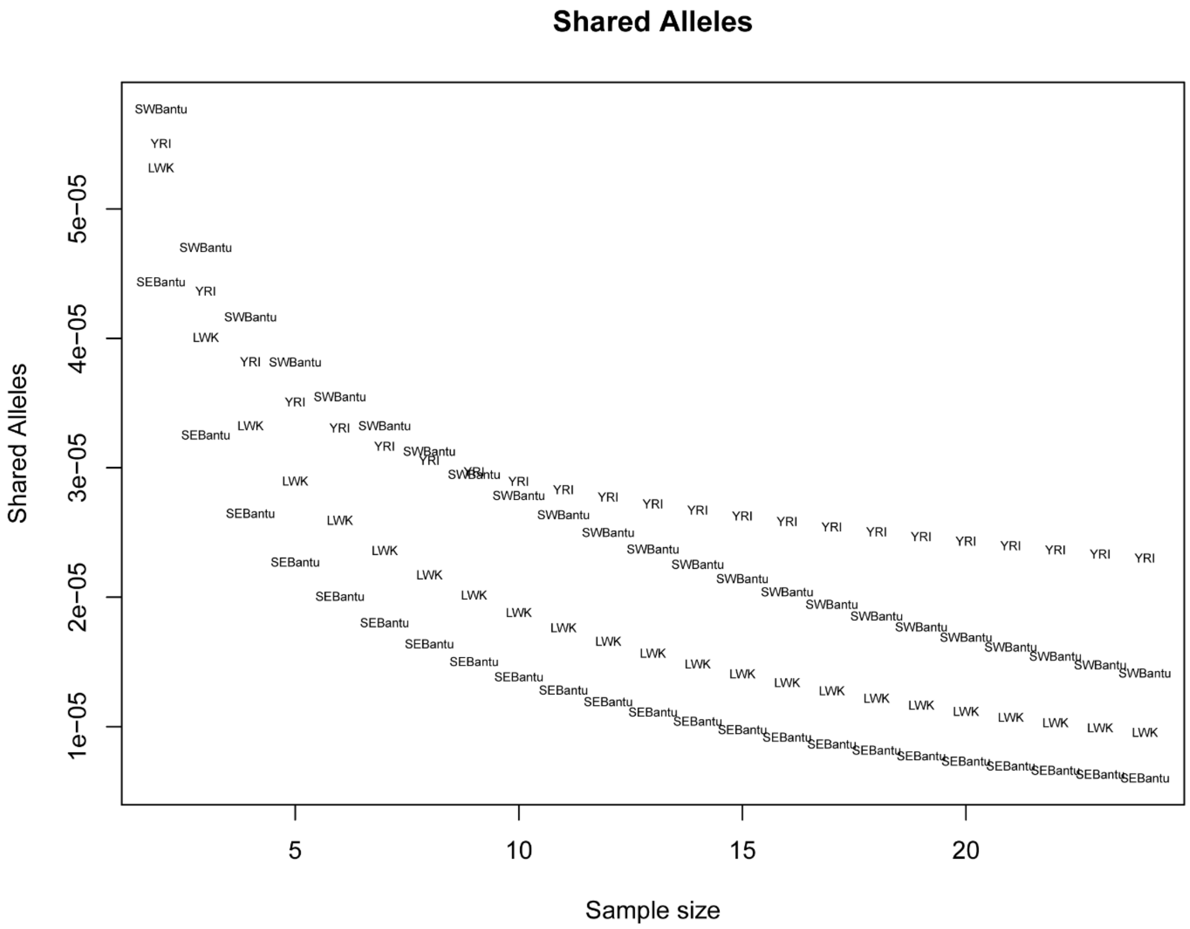


*Figure S7: Shared private alleles between the Afrikaner populations and populations with West-African ancestry.*

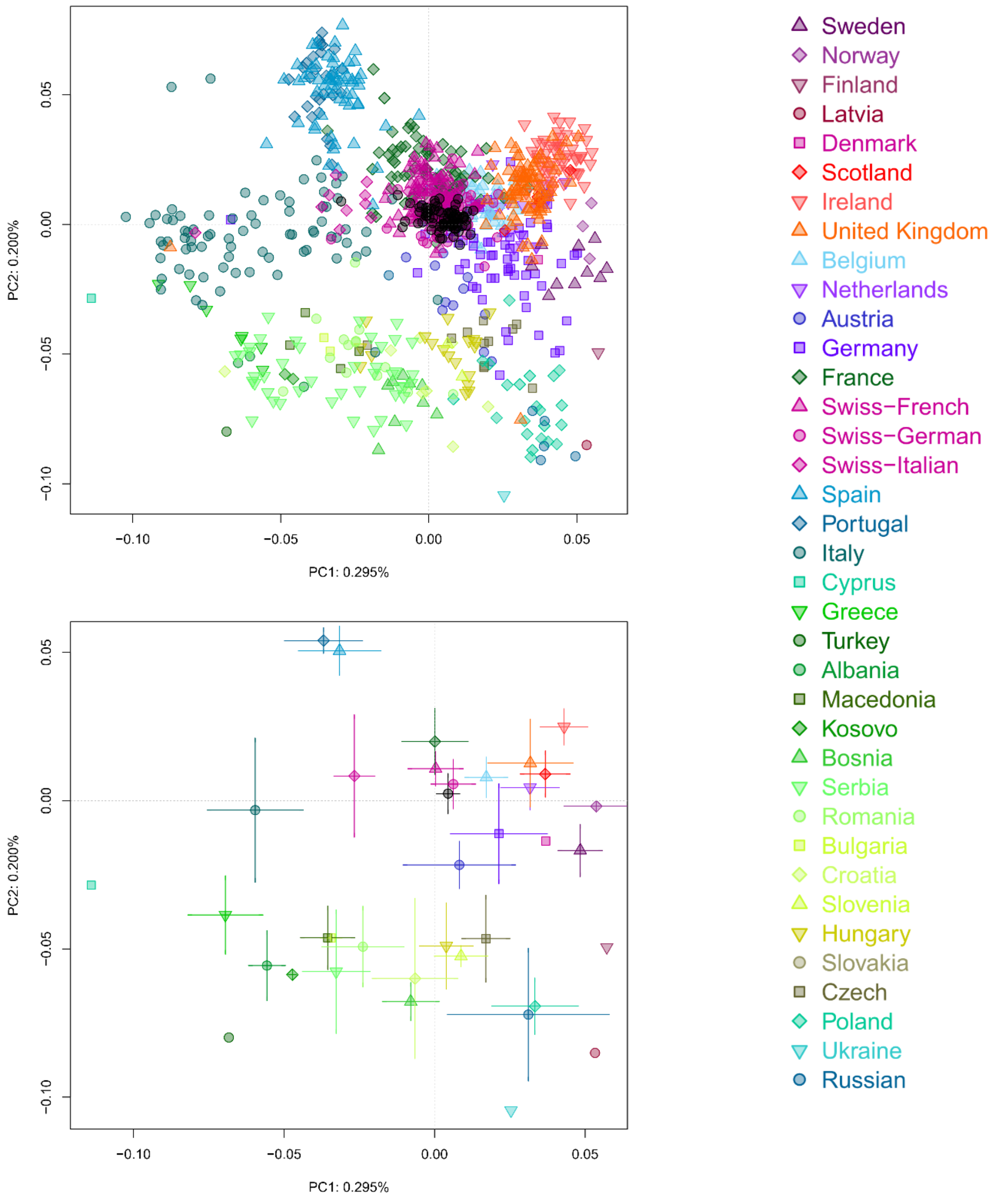


*Figure S8: A) Afrikaner individuals (black circles) projected on a PCA based on European genetic variation from the POPRES dataset. B) Population variation on PC 1 and 2 summarized as averages and standard deviations.*

A B

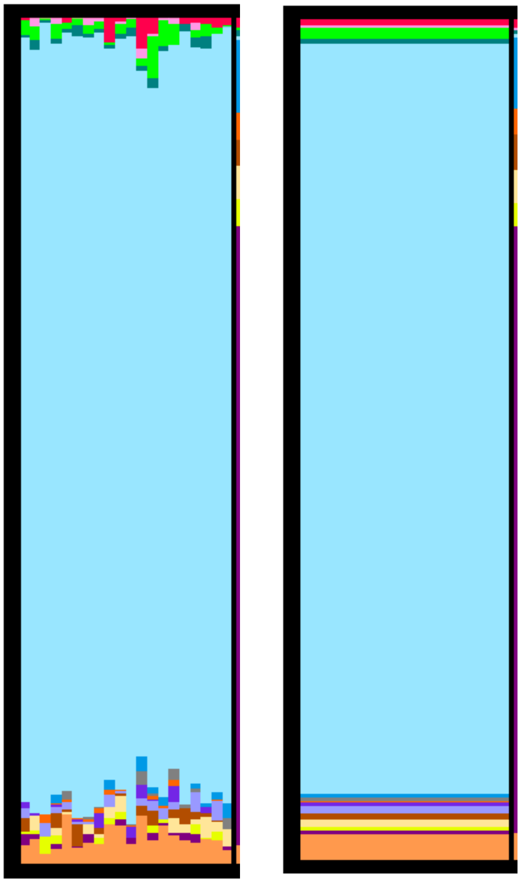

C D

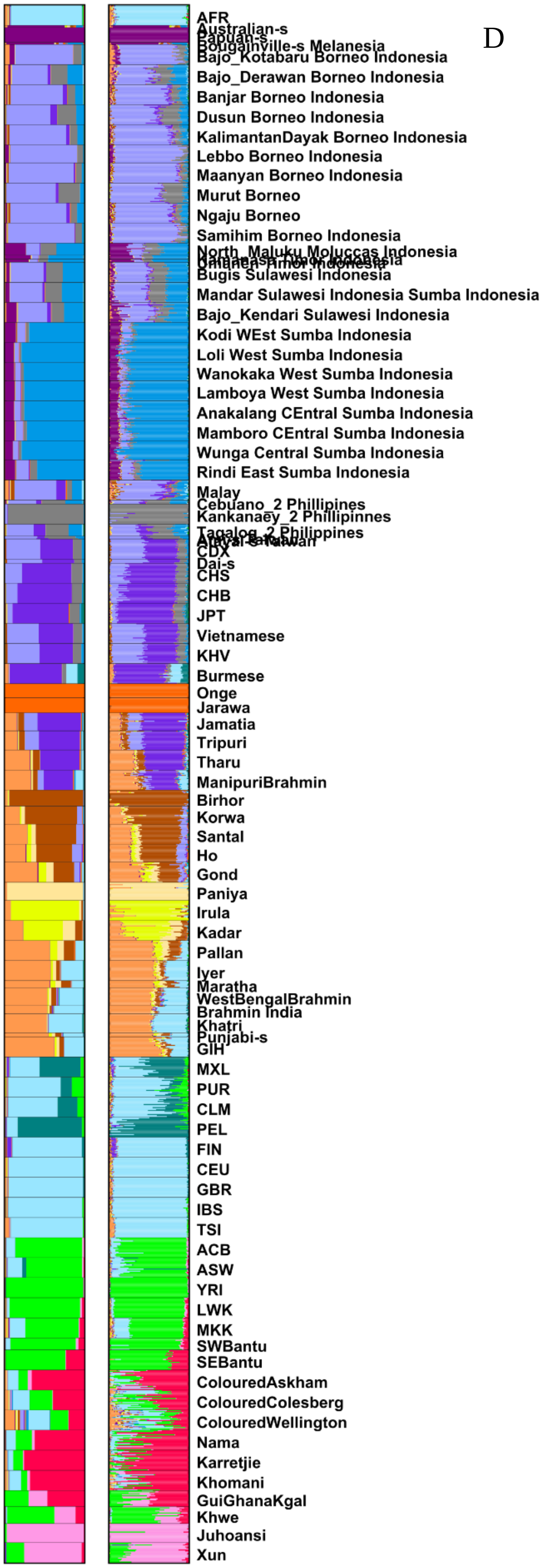


E

| Cluster Order | Color | Major cluster in: |
|---|---|---|
| 1 | red | Southern san |
| 2 | pink | Northern San |
| 3 | light_green | West African |
| 4 | sea_green | Native American |
| **5** | **light_blue** | **European** |
| 6 | blue | Sumba Indonesia |
| 7 | gray | Phillipines |
| 8 | dark_orange | Onge (Andaman) |
| 9 | purple | Japanese (East Asian) |
| 10 | light_purple | Borneo Indonesia, Malaysia |
| 11 | brown | India (Central, Austro_asiatic) |
| 12 | light_yellow | India (South, Dravidian) |
| 13 | yellow | India (South, Dravidian) |
| 14 | maroon | Australian and Papuan |
| 15 | orange | Indian (Upper Caste, Indo-European speaking) |

*Figure S9: Admixture analyses of the Asian extended dataset. Clustering at K=15. Panels A and B show a zoom-in of the AFR individuals, with A the individual clustering and B the population average clustering. Panels C and D show clustering at K=15 for the entire Asian extended dataset, with population average clustering (C) and individual clustering (D). Clusters were sorted so that non-Asian admixture are shown above the European component (light blue component) and Asian admixture below the European component at the bottom of the panel, according to the order shown in E.*

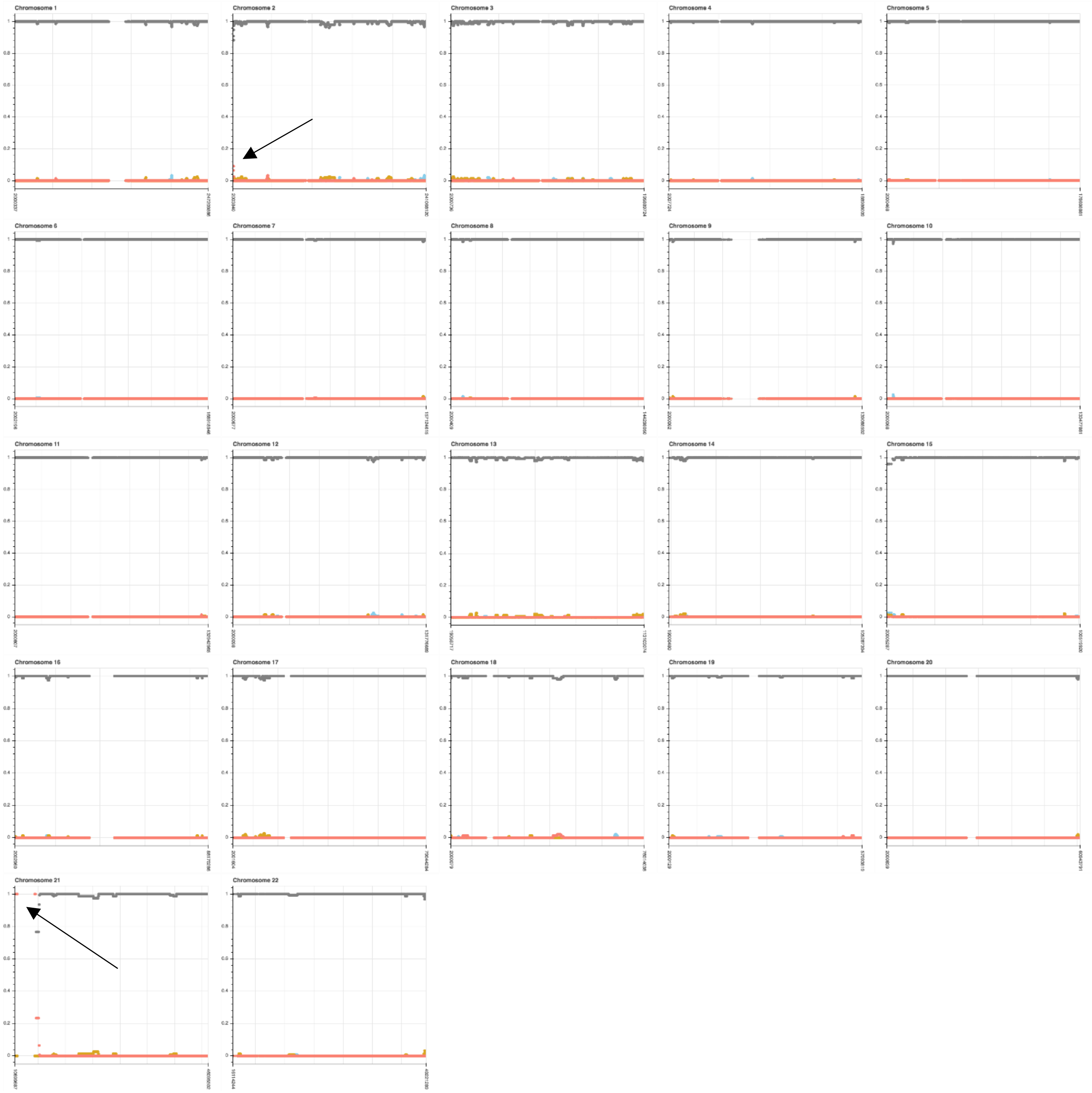


*Figure S10A: Local ancestry analyses. Different panes in the figure show different chromosomes. The x-axis indicate position along the genome and the y-axis the fraction of ancestry in percentage. Different colors indicate different ancestries, gray-European (CEU), blue-Asian (CDX), golden red- West African (YRI) and salmon - Khoe-San (combination of Khoe-San groups). Arrows indicate two positions discussed in the main text. (See Figure S10B below for zoom-in of chr 21)*

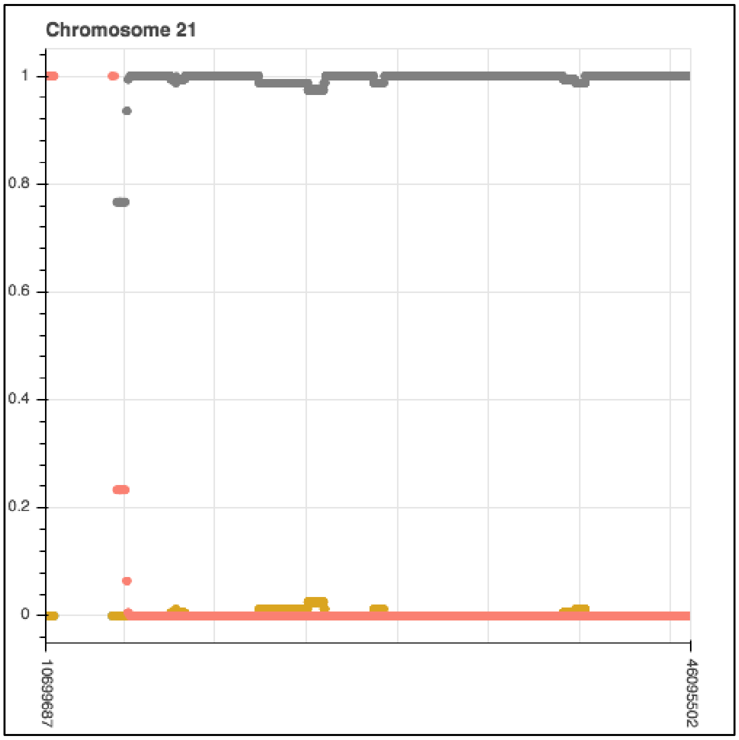


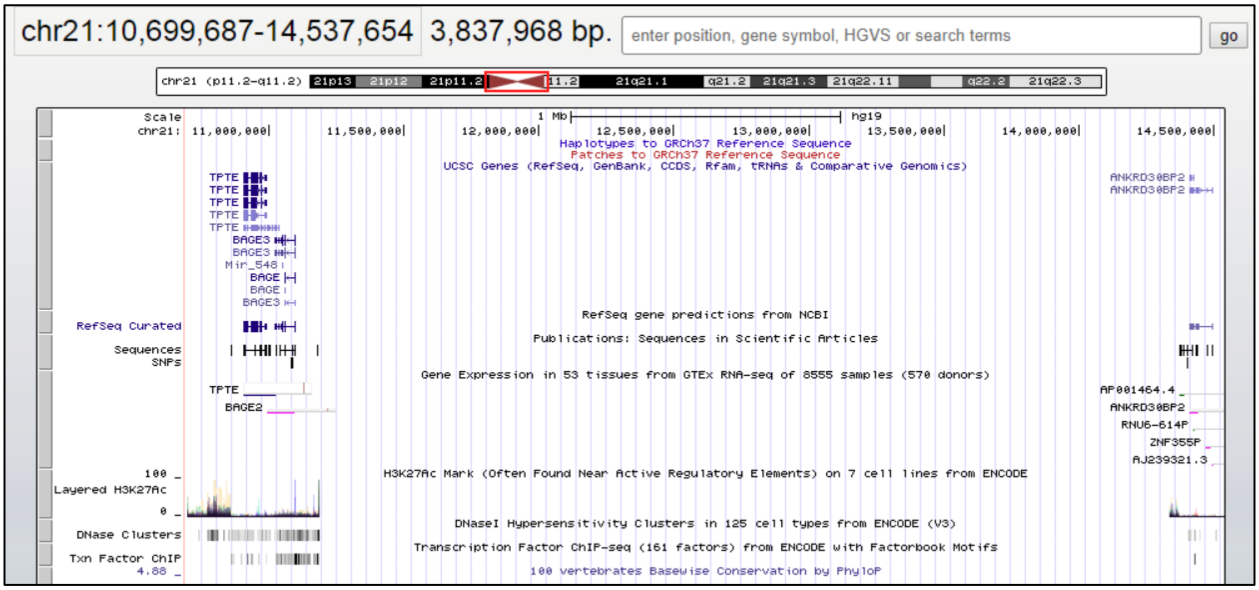


*Figure S10B: Local ancestry analyses. The top panel show a zoom-in of the local ancestry assignments on chromosome 21. The region between positions 10,699,687-14,537,654 seems to be 100% assigned to Khoe-San ancestry in 154 Afrikaner chromosomes. The panel at the bottom highlight the region in the UCSC genome browser, indicating the position of genes and the centromere.*

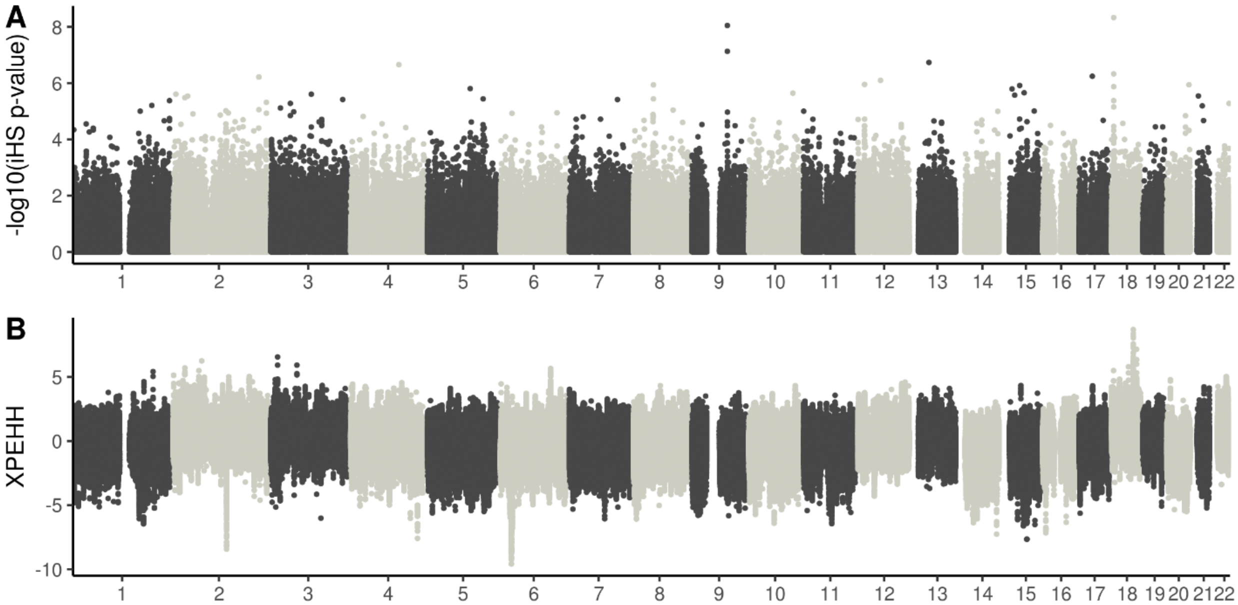


*Figure S11: Manhattan plots of selection scan results. A, iHS reslts and B, XP-EHH results. XP-EHH were calculated between AFR and CEU. Regions of extended homozygosity of AFR (compared to CEU) are indicated above the line and regions of extended homozygosity in CEU (compared to AFR) are indicated below the line. The X-axis indicate chromosomes and genomic positions and the Y-axis indicate –log10(p-value)*

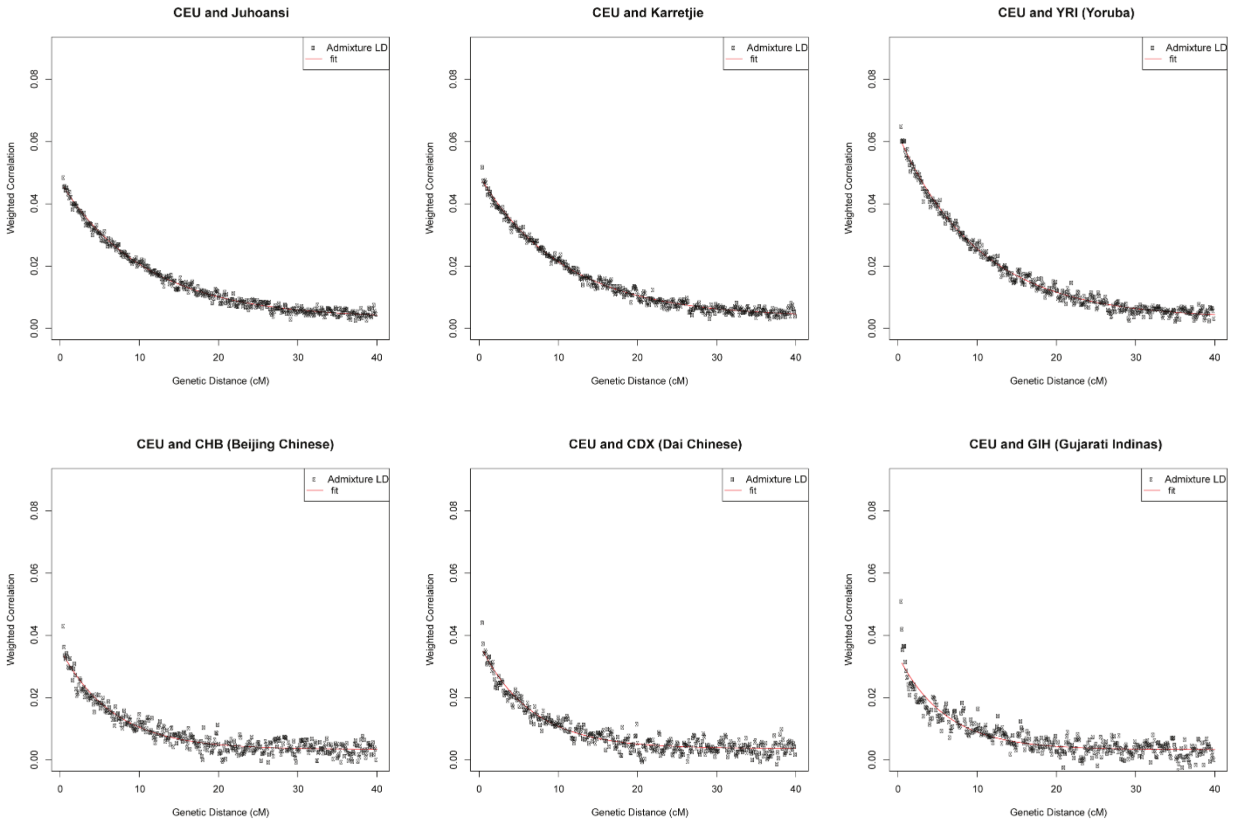


*Figure S12: LD-decay curves estimated with ROLLOFF.*

*Table S1: Admixture fractions of the Afrikaner individuals at K=6 (ADMIXTURE). Sorted by total fraction of Non-European ancestry.*

| ID | European | Total other Admixture | South-Asian | Khoe-San | East Asian | West/East African | Native American |
|---|---|---|---|---|---|---|---|
| 122 | 0.751 | 0.249 | 0.044 | 0.137 | 0.046 | 0.022 | 0.000 |
| 7 | 0.851 | 0.149 | 0.060 | 0.043 | 0.021 | 0.017 | 0.008 |
| 247 | 0.854 | 0.146 | 0.027 | 0.011 | 0.089 | 0.018 | 0.000 |
| 87 | 0.868 | 0.132 | 0.041 | 0.053 | 0.038 | 0.000 | 0.000 |
| 109 | 0.891 | 0.109 | 0.023 | 0.036 | 0.020 | 0.029 | 0.000 |
| 162 | 0.902 | 0.098 | 0.029 | 0.018 | 0.025 | 0.022 | 0.004 |
| 263 | 0.905 | 0.095 | 0.040 | 0.035 | 0.004 | 0.016 | 0.000 |
| 28 | 0.916 | 0.084 | 0.009 | 0.020 | 0.015 | 0.034 | 0.007 |
| 67 | 0.917 | 0.083 | 0.045 | 0.028 | 0.010 | 0.000 | 0.000 |
| 199 | 0.918 | 0.082 | 0.031 | 0.012 | 0.023 | 0.016 | 0.000 |
| 192 | 0.919 | 0.081 | 0.038 | 0.023 | 0.008 | 0.008 | 0.005 |
| 259 | 0.924 | 0.076 | 0.042 | 0.014 | 0.011 | 0.007 | 0.003 |
| 203 | 0.927 | 0.073 | 0.038 | 0.021 | 0.007 | 0.007 | 0.000 |
| 64 | 0.930 | 0.070 | 0.047 | 0.013 | 0.006 | 0.003 | 0.001 |
| 256 | 0.933 | 0.067 | 0.011 | 0.021 | 0.019 | 0.016 | 0.000 |
| 258 | 0.936 | 0.064 | 0.022 | 0.022 | 0.012 | 0.008 | 0.000 |
| 234 | 0.940 | 0.060 | 0.018 | 0.019 | 0.022 | 0.001 | 0.001 |
| 255 | 0.944 | 0.056 | 0.024 | 0.009 | 0.016 | 0.007 | 0.000 |
| 128 | 0.945 | 0.056 | 0.034 | 0.007 | 0.002 | 0.012 | 0.000 |
| 601 | 0.945 | 0.055 | 0.025 | 0.007 | 0.011 | 0.012 | 0.000 |
| 14 | 0.945 | 0.055 | 0.037 | 0.013 | 0.005 | 0.000 | 0.000 |
| 2 | 0.945 | 0.055 | 0.016 | 0.016 | 0.013 | 0.002 | 0.007 |
| 76 | 0.946 | 0.054 | 0.027 | 0.000 | 0.027 | 0.000 | 0.000 |
| 200 | 0.948 | 0.052 | 0.005 | 0.014 | 0.010 | 0.019 | 0.005 |
| 111 | 0.949 | 0.052 | 0.000 | 0.010 | 0.013 | 0.029 | 0.000 |
| 81 | 0.949 | 0.051 | 0.038 | 0.006 | 0.000 | 0.007 | 0.000 |
| 155 | 0.951 | 0.049 | 0.011 | 0.010 | 0.000 | 0.029 | 0.000 |
| 215 | 0.952 | 0.048 | 0.018 | 0.009 | 0.011 | 0.011 | 0.000 |
| 51 | 0.952 | 0.048 | 0.011 | 0.007 | 0.001 | 0.028 | 0.002 |
| 238 | 0.954 | 0.046 | 0.019 | 0.005 | 0.000 | 0.018 | 0.004 |
| 241 | 0.956 | 0.044 | 0.004 | 0.013 | 0.005 | 0.022 | 0.000 |
| 46 | 0.957 | 0.043 | 0.021 | 0.007 | 0.008 | 0.008 | 0.000 |
| 261 | 0.957 | 0.043 | 0.021 | 0.014 | 0.007 | 0.000 | 0.000 |
| 605 | 0.958 | 0.042 | 0.011 | 0.012 | 0.005 | 0.013 | 0.001 |
| 604 | 0.959 | 0.041 | 0.003 | 0.016 | 0.018 | 0.004 | 0.000 |
| 154 | 0.961 | 0.040 | 0.014 | 0.009 | 0.016 | 0.000 | 0.000 |
| 260 | 0.961 | 0.039 | 0.025 | 0.008 | 0.000 | 0.005 | 0.001 |
| 229 | 0.961 | 0.039 | 0.024 | 0.006 | 0.001 | 0.009 | 0.000 |
| 22 | 0.961 | 0.039 | 0.012 | 0.018 | 0.000 | 0.006 | 0.002 |
| 49 | 0.961 | 0.039 | 0.023 | 0.007 | 0.005 | 0.005 | 0.000 |
| 89 | 0.962 | 0.038 | 0.004 | 0.020 | 0.008 | 0.007 | 0.000 |

| | | | | | | | |
|---|---|---|---|---|---|---|---|
| 120 | 0.963 | 0.037 | 0.024 | 0.002 | 0.011 | 0.000 | 0.000 |
| 262 | 0.964 | 0.036 | 0.000 | 0.020 | 0.012 | 0.003 | 0.001 |
| 1 | 0.966 | 0.034 | 0.012 | 0.013 | 0.006 | 0.004 | 0.000 |
| 194 | 0.967 | 0.033 | 0.014 | 0.008 | 0.002 | 0.010 | 0.000 |
| 218 | 0.967 | 0.033 | 0.017 | 0.009 | 0.002 | 0.000 | 0.005 |
| 21 | 0.967 | 0.033 | 0.012 | 0.005 | 0.015 | 0.000 | 0.001 |
| 35 | 0.969 | 0.031 | 0.021 | 0.010 | 0.000 | 0.000 | 0.000 |
| 144 | 0.970 | 0.030 | 0.000 | 0.007 | 0.013 | 0.009 | 0.000 |
| 264 | 0.970 | 0.030 | 0.013 | 0.007 | 0.006 | 0.003 | 0.000 |
| 129 | 0.971 | 0.029 | 0.010 | 0.008 | 0.002 | 0.006 | 0.004 |
| 138 | 0.972 | 0.028 | 0.000 | 0.010 | 0.003 | 0.015 | 0.000 |
| 97 | 0.973 | 0.027 | 0.008 | 0.001 | 0.017 | 0.000 | 0.002 |
| 153 | 0.973 | 0.027 | 0.014 | 0.006 | 0.004 | 0.000 | 0.003 |
| 225 | 0.973 | 0.027 | 0.004 | 0.007 | 0.000 | 0.016 | 0.000 |
| 160 | 0.974 | 0.026 | 0.009 | 0.006 | 0.000 | 0.012 | 0.000 |
| 18 | 0.974 | 0.026 | 0.016 | 0.006 | 0.004 | 0.000 | 0.000 |
| 24 | 0.976 | 0.025 | 0.000 | 0.013 | 0.001 | 0.010 | 0.000 |
| 201 | 0.977 | 0.023 | 0.000 | 0.008 | 0.001 | 0.014 | 0.000 |
| 267 | 0.977 | 0.023 | 0.012 | 0.005 | 0.006 | 0.000 | 0.000 |
| 254 | 0.977 | 0.023 | 0.004 | 0.011 | 0.000 | 0.008 | 0.000 |
| 19 | 0.977 | 0.023 | 0.020 | 0.000 | 0.000 | 0.000 | 0.003 |
| 189 | 0.978 | 0.023 | 0.003 | 0.007 | 0.000 | 0.013 | 0.000 |
| 226 | 0.978 | 0.022 | 0.014 | 0.008 | 0.001 | 0.000 | 0.000 |
| 178 | 0.978 | 0.022 | 0.015 | 0.002 | 0.005 | 0.000 | 0.000 |
| 56 | 0.978 | 0.022 | 0.004 | 0.011 | 0.003 | 0.000 | 0.004 |
| 219 | 0.979 | 0.021 | 0.019 | 0.002 | 0.000 | 0.000 | 0.000 |
| 180 | 0.980 | 0.020 | 0.010 | 0.005 | 0.000 | 0.005 | 0.000 |
| 94 | 0.981 | 0.020 | 0.000 | 0.003 | 0.007 | 0.009 | 0.000 |
| 57 | 0.983 | 0.017 | 0.012 | 0.000 | 0.004 | 0.000 | 0.000 |
| 150 | 0.984 | 0.017 | 0.000 | 0.007 | 0.009 | 0.000 | 0.001 |
| 265 | 0.990 | 0.010 | 0.000 | 0.004 | 0.000 | 0.000 | 0.005 |
| 137 | 0.991 | 0.009 | 0.000 | 0.003 | 0.000 | 0.005 | 0.000 |
| 257 | 0.995 | 0.005 | 0.000 | 0.005 | 0.000 | 0.000 | 0.000 |
| 86 | 0.997 | 0.003 | 0.000 | 0.000 | 0.001 | 0.000 | 0.002 |
| 53 | 0.997 | 0.003 | 0.002 | 0.000 | 0.000 | 0.000 | 0.001 |
| 31 | 1.000 | 0.000 | 0.000 | 0.000 | 0.000 | 0.000 | 0.000 |
| **Mean** | **0.953** | **0.047** | **0.017** | **0.013** | **0.009** | **0.008** | **0.001** |
| **SD** | **0.038** | **0.038** | **0.014** | **0.017** | **0.013** | **0.009** | **0.002** |
| **Median** | **0.961** | **0.039** | **0.014** | **0.009** | **0.005** | **0.006** | **0.000** |
| **MIN** | **0.751** | **0.000** | **0.000** | **0.000** | **0.000** | **0.000** | **0.000** |
| **MAX** | **1.000** | **0.249** | **0.060** | **0.137** | **0.089** | **0.034** | **0.008** |
| **Count if 0** | **0** | **1** | **12** | **6** | **17** | **26** | **51** |

*Table S2: Admixture fractions of the Afrikaner individuals at K=9 (ADMIXTURE). Sorted by total fraction of non-European ancestry.*

| ID | North Eur | South Eur | Total other Admixture | SE Asian | Khoe-San | West Afr | Chinese | East Afr | Japanese | Native American |
|---|---|---|---|---|---|---|---|---|---|---|
| 122 | 0.403 | 0.354 | 0.243 | 0.040 | 0.133 | 0.000 | 0.039 | 0.031 | 0.000 | 0.000 |
| 7 | 0.466 | 0.396 | 0.139 | 0.061 | 0.040 | 0.005 | 0.016 | 0.017 | 0.000 | 0.000 |
| 247 | 0.482 | 0.381 | 0.137 | 0.029 | 0.009 | 0.018 | 0.080 | 0.000 | 0.000 | 0.000 |
| 87 | 0.493 | 0.384 | 0.123 | 0.042 | 0.053 | 0.000 | 0.028 | 0.000 | 0.000 | 0.000 |
| 109 | 0.439 | 0.461 | 0.100 | 0.024 | 0.034 | 0.022 | 0.010 | 0.011 | 0.000 | 0.000 |
| 162 | 0.427 | 0.484 | 0.089 | 0.030 | 0.016 | 0.021 | 0.020 | 0.001 | 0.000 | 0.000 |
| 263 | 0.447 | 0.468 | 0.085 | 0.036 | 0.034 | 0.015 | 0.000 | 0.000 | 0.000 | 0.000 |
| 67 | 0.467 | 0.458 | 0.075 | 0.046 | 0.027 | 0.000 | 0.002 | 0.000 | 0.000 | 0.000 |
| 28 | 0.443 | 0.483 | 0.074 | 0.010 | 0.017 | 0.028 | 0.009 | 0.009 | 0.000 | 0.001 |
| 199 | 0.442 | 0.485 | 0.073 | 0.031 | 0.010 | 0.016 | 0.016 | 0.000 | 0.000 | 0.000 |
| 192 | 0.469 | 0.460 | 0.071 | 0.040 | 0.021 | 0.000 | 0.000 | 0.009 | 0.000 | 0.000 |
| 259 | 0.490 | 0.444 | 0.067 | 0.042 | 0.013 | 0.000 | 0.004 | 0.008 | 0.000 | 0.000 |
| 203 | 0.464 | 0.472 | 0.064 | 0.038 | 0.020 | 0.006 | 0.000 | 0.000 | 0.000 | 0.000 |
| 64 | 0.495 | 0.444 | 0.061 | 0.046 | 0.010 | 0.000 | 0.000 | 0.005 | 0.000 | 0.000 |
| 256 | 0.509 | 0.433 | 0.058 | 0.011 | 0.018 | 0.001 | 0.010 | 0.020 | 0.000 | 0.000 |
| 258 | 0.459 | 0.485 | 0.056 | 0.022 | 0.020 | 0.000 | 0.005 | 0.009 | 0.000 | 0.000 |
| 234 | 0.484 | 0.464 | 0.052 | 0.019 | 0.017 | 0.000 | 0.013 | 0.003 | 0.000 | 0.000 |
| 128 | 0.411 | 0.541 | 0.048 | 0.028 | 0.003 | 0.000 | 0.000 | 0.016 | 0.000 | 0.000 |
| 255 | 0.488 | 0.464 | 0.048 | 0.024 | 0.008 | 0.006 | 0.009 | 0.000 | 0.000 | 0.000 |
| 76 | 0.485 | 0.468 | 0.047 | 0.030 | 0.000 | 0.000 | 0.011 | 0.000 | 0.006 | 0.000 |
| 601 | 0.482 | 0.471 | 0.047 | 0.027 | 0.005 | 0.011 | 0.003 | 0.000 | 0.000 | 0.000 |
| 14 | 0.490 | 0.464 | 0.045 | 0.034 | 0.011 | 0.000 | 0.000 | 0.000 | 0.000 | 0.000 |
| 2 | 0.513 | 0.444 | 0.043 | 0.020 | 0.016 | 0.000 | 0.008 | 0.000 | 0.000 | 0.000 |
| 111 | 0.484 | 0.473 | 0.043 | 0.000 | 0.006 | 0.019 | 0.004 | 0.014 | 0.000 | 0.000 |
| 81 | 0.506 | 0.452 | 0.042 | 0.032 | 0.004 | 0.005 | 0.000 | 0.001 | 0.000 | 0.000 |
| 200 | 0.504 | 0.455 | 0.041 | 0.007 | 0.012 | 0.019 | 0.003 | 0.000 | 0.000 | 0.000 |
| 215 | 0.499 | 0.461 | 0.040 | 0.019 | 0.007 | 0.003 | 0.002 | 0.010 | 0.000 | 0.000 |
| 155 | 0.493 | 0.468 | 0.039 | 0.003 | 0.008 | 0.028 | 0.000 | 0.000 | 0.000 | 0.000 |
| 51 | 0.499 | 0.464 | 0.038 | 0.005 | 0.005 | 0.028 | 0.000 | 0.000 | 0.000 | 0.000 |
| 46 | 0.491 | 0.473 | 0.036 | 0.022 | 0.005 | 0.000 | 0.000 | 0.009 | 0.000 | 0.000 |
| 238 | 0.472 | 0.493 | 0.036 | 0.015 | 0.003 | 0.018 | 0.000 | 0.000 | 0.000 | 0.000 |
| 241 | 0.485 | 0.480 | 0.035 | 0.001 | 0.010 | 0.016 | 0.000 | 0.009 | 0.000 | 0.000 |
| 261 | 0.441 | 0.524 | 0.034 | 0.021 | 0.013 | 0.000 | 0.001 | 0.000 | 0.000 | 0.000 |
| 605 | 0.453 | 0.513 | 0.034 | 0.009 | 0.010 | 0.006 | 0.000 | 0.008 | 0.000 | 0.000 |
| 604 | 0.509 | 0.460 | 0.032 | 0.005 | 0.015 | 0.003 | 0.010 | 0.000 | 0.000 | 0.000 |
| 229 | 0.531 | 0.437 | 0.031 | 0.017 | 0.003 | 0.000 | 0.000 | 0.012 | 0.000 | 0.000 |
| 154 | 0.476 | 0.494 | 0.031 | 0.012 | 0.007 | 0.000 | 0.010 | 0.001 | 0.000 | 0.000 |
| 49 | 0.469 | 0.500 | 0.031 | 0.021 | 0.005 | 0.000 | 0.000 | 0.005 | 0.000 | 0.000 |
| 260 | 0.521 | 0.450 | 0.030 | 0.018 | 0.007 | 0.000 | 0.000 | 0.004 | 0.000 | 0.000 |
| 120 | 0.471 | 0.500 | 0.029 | 0.024 | 0.000 | 0.000 | 0.005 | 0.000 | 0.000 | 0.000 |
| 89 | 0.521 | 0.450 | 0.029 | 0.002 | 0.017 | 0.000 | 0.000 | 0.010 | 0.000 | 0.000 |
| 22 | 0.490 | 0.481 | 0.028 | 0.004 | 0.016 | 0.000 | 0.000 | 0.008 | 0.000 | 0.000 |

| | | | | | | | | | | |
|---|---|---|---|---|---|---|---|---|---|---|
| 262 | 0.492 | 0.482 | 0.027 | 0.000 | 0.018 | 0.000 | 0.005 | 0.003 | 0.000 | 0.000 |
| 194 | 0.514 | 0.461 | 0.025 | 0.010 | 0.006 | 0.007 | 0.000 | 0.003 | 0.000 | 0.000 |
| 1 | 0.479 | 0.496 | 0.025 | 0.009 | 0.011 | 0.000 | 0.000 | 0.005 | 0.000 | 0.000 |
| 21 | 0.555 | 0.422 | 0.024 | 0.014 | 0.003 | 0.000 | 0.007 | 0.000 | 0.000 | 0.000 |
| 138 | 0.450 | 0.528 | 0.023 | 0.000 | 0.009 | 0.013 | 0.000 | 0.001 | 0.000 | 0.000 |
| 218 | 0.472 | 0.505 | 0.023 | 0.015 | 0.008 | 0.000 | 0.000 | 0.000 | 0.000 | 0.000 |
| 35 | 0.515 | 0.463 | 0.022 | 0.013 | 0.009 | 0.000 | 0.000 | 0.000 | 0.000 | 0.000 |
| 24 | 0.537 | 0.442 | 0.021 | 0.000 | 0.011 | 0.004 | 0.000 | 0.006 | 0.000 | 0.000 |
| 264 | 0.469 | 0.511 | 0.021 | 0.013 | 0.006 | 0.002 | 0.000 | 0.000 | 0.000 | 0.000 |
| 225 | 0.515 | 0.465 | 0.020 | 0.000 | 0.005 | 0.015 | 0.000 | 0.000 | 0.000 | 0.000 |
| 201 | 0.467 | 0.513 | 0.020 | 0.000 | 0.007 | 0.012 | 0.000 | 0.001 | 0.000 | 0.000 |
| 144 | 0.496 | 0.484 | 0.020 | 0.001 | 0.007 | 0.005 | 0.004 | 0.003 | 0.000 | 0.000 |
| 129 | 0.509 | 0.472 | 0.020 | 0.006 | 0.005 | 0.000 | 0.000 | 0.009 | 0.000 | 0.000 |
| 189 | 0.512 | 0.469 | 0.020 | 0.000 | 0.003 | 0.000 | 0.000 | 0.016 | 0.000 | 0.000 |
| 18 | 0.490 | 0.492 | 0.018 | 0.014 | 0.004 | 0.000 | 0.000 | 0.000 | 0.000 | 0.000 |
| 153 | 0.448 | 0.534 | 0.018 | 0.013 | 0.005 | 0.000 | 0.000 | 0.000 | 0.000 | 0.000 |
| 160 | 0.502 | 0.481 | 0.018 | 0.000 | 0.003 | 0.002 | 0.000 | 0.013 | 0.000 | 0.000 |
| 97 | 0.500 | 0.483 | 0.017 | 0.008 | 0.000 | 0.000 | 0.010 | 0.000 | 0.000 | 0.000 |
| 254 | 0.502 | 0.482 | 0.016 | 0.000 | 0.010 | 0.006 | 0.000 | 0.000 | 0.000 | 0.000 |
| 178 | 0.500 | 0.487 | 0.014 | 0.014 | 0.000 | 0.000 | 0.000 | 0.000 | 0.000 | 0.000 |
| 19 | 0.464 | 0.522 | 0.013 | 0.013 | 0.000 | 0.000 | 0.000 | 0.000 | 0.000 | 0.000 |
| 267 | 0.501 | 0.486 | 0.013 | 0.011 | 0.003 | 0.000 | 0.000 | 0.000 | 0.000 | 0.000 |
| 226 | 0.474 | 0.514 | 0.013 | 0.007 | 0.005 | 0.000 | 0.000 | 0.000 | 0.000 | 0.000 |
| 56 | 0.479 | 0.509 | 0.012 | 0.001 | 0.010 | 0.000 | 0.000 | 0.000 | 0.000 | 0.001 |
| 219 | 0.472 | 0.517 | 0.012 | 0.012 | 0.000 | 0.000 | 0.000 | 0.000 | 0.000 | 0.000 |
| 180 | 0.498 | 0.492 | 0.010 | 0.001 | 0.002 | 0.000 | 0.000 | 0.007 | 0.000 | 0.000 |
| 94 | 0.474 | 0.517 | 0.009 | 0.000 | 0.003 | 0.006 | 0.001 | 0.000 | 0.000 | 0.000 |
| 57 | 0.445 | 0.547 | 0.008 | 0.008 | 0.000 | 0.000 | 0.000 | 0.000 | 0.000 | 0.000 |
| 150 | 0.500 | 0.492 | 0.008 | 0.002 | 0.006 | 0.000 | 0.000 | 0.000 | 0.000 | 0.000 |
| 137 | 0.514 | 0.481 | 0.005 | 0.000 | 0.003 | 0.000 | 0.000 | 0.002 | 0.000 | 0.000 |
| 265 | 0.450 | 0.548 | 0.002 | 0.000 | 0.002 | 0.000 | 0.000 | 0.000 | 0.000 | 0.000 |
| 257 | 0.519 | 0.479 | 0.002 | 0.000 | 0.002 | 0.000 | 0.000 | 0.000 | 0.000 | 0.000 |
| 31 | 0.479 | 0.521 | 0.000 | 0.000 | 0.000 | 0.000 | 0.000 | 0.000 | 0.000 | 0.000 |
| 53 | 0.497 | 0.503 | 0.000 | 0.000 | 0.000 | 0.000 | 0.000 | 0.000 | 0.000 | 0.000 |
| 86 | 0.490 | 0.510 | 0.000 | 0.000 | 0.000 | 0.000 | 0.000 | 0.000 | 0.000 | 0.000 |
| **Mean** | **0.483599** | **0.477206** | **0.039162** | **0.014923** | **0.011065** | **0.004751** | **0.004471** | **0.003847** | **0.000083** | **0.000022** |
| **SD** | **0.027902** | **0.035821** | **0.037591** | **0.014403** | **0.017047** | **0.007833** | **0.011060** | **0.005963** | **0.000729** | **0.000142** |
| **Median** | **0.488300** | **0.480700** | **0.029500** | **0.012000** | **0.006800** | **0.000000** | **0.000000** | **0.000000** | **0.000000** | **0.000000** |
| **MIN** | **0.403100** | **0.354400** | **0.000000** | **0.000000** | **0.000000** | **0.000000** | **0.000000** | **0.000000** | **0.000000** | **0.000000** |
| **MAX** | **0.554600** | **0.548400** | **0.242600** | **0.060500** | **0.133200** | **0.028400** | **0.080200** | **0.030800** | **0.006400** | **0.001100** |
| **Count if 0** | **0** | **0** | **3** | **16** | **9** | **46** | **47** | **40** | **76** | **75** |

*Table S3: Top 5 selection scan peaks detected with iHS and XP-EHH scans*

| **Peak** | **Position information** (chr:hg19 chr position) | **Underlying or nearby genes*** | **Potential function of underlying/nearby genes*** |
|---|---|---|---|
| iHS-1 | 18:7486106 | No underlying genes, Closest gene: PTPRM (top SNP in peak is 82 kb upstream from gene) | PTPRM: member of the protein tyrosine phosphatase (PTP) family. PTPs are known to be signaling molecules that regulate a variety of cellular processes including cell growth, differentiation, mitotic cycle, and oncogenic transformation |
| iHS-2 | 9:90898783 | No underlying genes, Closest gene: SPIN1 (top SNP in peak is 143 kb upstream from gene) | Homo sapiens spindlin 1 (SPIN1), mRNA. No known function |
| iHS-3 | 13:45630008 | No underlying genes, Closest gene: KIAA1704 (top SNP in peak is 41 kb downstream from gene) | Homo sapiens KIAA1704 (KIAA1704), mRNA. No known function |
| iHS-4 | 4:123774065 | Underlying gene: FGF2 | Homo sapiens fibroblast growth factor 2 (FGF2). The protein encoded by this gene is a member of the fibroblast growth factor (FGF) family. FGF family members bind heparin and possess broad mitogenic and angiogenic activities. This protein has been implicated in diverse biological processes, such as limb and nervous system development, wound healing, and tumor growth. Positive Disease Associations: Cholesterol, LDL [PubMed 17903299] |
| iHS-5 | 17:36185998 | No underlying genes, Closest genes are:<br><br>LOC284100 (top SNP in peak is 17 kb downstream from gene)<br><br>HNF1B: (top SNP in peak is 139 kb upstream from gene) | LOC284100: Homo sapiens tyrosine 3-monooxygenase/tryptophan 5-monooxygenase activation protein, epsilon polypeptide pseudogene (LOC284100), non-coding RNA. No known function<br><br>HNF1B: Homo sapiens HNF1 homeobox B. This gene encodes a member of the homeodomain-containing superfamily of transcription factors. The protein binds to DNA as either a homodimer, or a heterodimer with the related protein hepatocyte nuclear factor 1-alpha. The gene has been shown to function in nephron development, and regulates development of the embryonic pancreas. Mutations in this gene result in renal cysts and diabetes syndrome and noninsulin-dependent diabetes mellitus, and expression of this gene is altered in some types of cancer. |
| XP-EHH-1 | 18:57128753 | Underlying gene: CCBE1 | Homo sapiens collagen and calcium binding EGF domains 1 (CCBE1). This gene is thought to function in extracellular matrix remodeling and migration. It is predominantly expressed in the ovary, but down regulated in ovarian cancer cell lines and primary carcinomas, suggesting its role as a tumour suppressor. Mutations in this gene have been associated with Hennekam lymphangiectasia-lymphedema syndrome, a generalized lymphatic dysplasia in humans. |

| | | | |
|---|---|---|---|
| | | | Positive Disease Associations: Alcoholism, Apolipoproteins B, Arteries, Blood Pressure Determination, Body Mass Index, Cell Adhesion Molecules, Cholesterol, Insulin, Rheumatoid Arthritis |
| XP-EHH-2 | 3:18609478 | No underlying genes, Closest gene:<br><br>SATB1 (top SNP in peak is 220 kb upstream from gene) | Homo sapiens SATB homeobox 1 (SATB1). This gene encodes a matrix protein which binds nuclear matrix and scaffold-associating DNAs through a unique nuclear architecture. The protein recruits chromatin-remodeling factors in order to regulate chromatin structure and gene expression. Crucial silencing factor contributing to the initiation of X inactivation mediated by Xist RNA that occurs during embryogenesis and in lymphoma. Positive Disease Associations: Alcoholism, Arteries, Cholesterol, HDL, Crohn Disease, Eosinophils, Inflammatory Bowel Diseases, Insulin, Insulin Resistance, Psoriasis |
| XP-EHH-3 | 2:74122223 | Underlying gene: ACTG2 | Homo sapiens actin, gamma 2, smooth muscle, enteric (ACTG2). Actins are highly conserved proteins that are involved in various types of cell motility and in the maintenance of the cytoskeleton. This gene encodes actin gamma 2; a smooth muscle actin found in enteric tissues. Diseases sorted by gene-association score: visceral myopathy* (1599), chronic intestinal pseudoobstruction* (400), myopathic intestinal pseudoobstruction* (350), actg2 (20), intestinal pseudo-obstruction (13), peyronie's disease (8), myopathy (3) |
| XP-EHH-4 | 3:67456973 | Underlying gene: SUCLG2 | Homo sapiens succinate-CoA ligase, GDP-forming, beta subunit (SUCLG2), nuclear gene encoding mitochondrial protein. This gene encodes a GTP-specific beta subunit of succinyl-CoA synthetase. Succinyl-CoA synthetase catalyzes the reversible reaction involving the formation of succinyl-CoA and succinate. Positive Disease Associations: Astigmatism , Glucose , Lipoproteins, VLDL |
| XP-EHH-5 | 6:128989302 | No underlying genes, Closest gene:<br><br>LAMA2 (top SNP in peak is 215 kb upstream from gene) | Homo sapiens laminin, alpha 2 (LAMA2). Laminin, an extracellular protein, is a major component of the basement membrane. It is thought to mediate the attachment, migration, and organization of cells into tissues during embryonic development by interacting with other extracellular matrix components. Mutations in this gene have been identified as the cause of congenital merosin-deficient muscular dystrophy. Positive Disease Associations: Amyotrophic Lateral Sclerosis, Body Mass Index, Body Weight, Cholesterol, HDL, Echocardiography, Exercise Test, leprosy, tuberculoid type of leprosy, Uric Acid |

*Underlying/nearby gene information and gene functions were obtained from UCSC genome browser (https://genome.ucsc.edu/)

*Table S4: Admixture LD decay estimate of admixture times into the Afrikaner population*

| Parental 1 | Parental 2 | Admixed | Generations | SE | Time +/- SE (29 y/gen) |
|---|---|---|---|---|---|
| CEU | Juhoansi | AFR | 9.222 | 0.434 | 267.4 +/- 12.6 |
| CEU | Karretjie | AFR | 9.383 | 0.465 | 272.1 +/- 13.5 |
| CEU | Khomani | AFR | 9.464 | 0.474 | 274.5 +/- 13.7 |
| CEU | YRI | AFR | 9.703 | 0.449 | 281.4 +/- 13.0 |
| CEU | CDX | AFR | 15.332 | 2.648 | 444.6 +/- 76.8 |
| CEU | CHB | AFR | 14.929 | 2.665 | 432.9 +/- 77.3 |
| CEU | GIH | AFR | 16.522 | 2.237 | 479.1 +/- 64.9 |

*Table S5: Average inbreeding coefficients per population, sorted in ascending order.*

| | |
|---|---|
| ASW | -0.00088 |
| ACB | -0.00054 |
| YRI | 2.60E-05 |
| MKK | 4.77E-05 |
| ColouredWellington | 0.000101 |
| SWBantu | 0.000171 |
| ColouredColesberg | 0.000233 |
| SEBantu | 0.000337 |
| LWK | 0.000365 |
| Khwe | 0.000629 |
| Nama | 0.000807 |
| ColouredAskham | 0.001006 |
| Khomani | 0.001032 |
| Karretjie | 0.001297 |
| GuiGhanaKgal | 0.001345 |
| Xun | 0.001476 |
| Juhoansi | 0.002185 |
| PUR | 0.002335 |
| MXL | 0.00284 |
| CLM | 0.003061 |
| AFR | 0.004128 |
| CEU | 0.004533 |
| TSI | 0.004603 |
| IBS | 0.004625 |
| GIH | 0.004718 |
| GBR | 0.004832 |
| FIN | 0.004967 |
| PEL | 0.005641 |
| KHV | 0.007984 |
| CHB | 0.007996 |
| JPT | 0.008152 |
| CHS | 0.008183 |
| CDX | 0.00836 |

### Supplementary Text

#### Supplementary Note 1

This supplementary note discusses the populations from which the Afrikaner population arose, it summarizes genealogical information of admixture and presents genetic information of admixture.

#### Population figures

*1. Khoekhoe and San*

The population of Khoekhoe and San, hereafter referred to as Khoe-San is estimated to have numbered 20,000 overall and 5,000 people in the western Cape in 1652 (Elphick and Malherbe 1989). Annexation of Khoe-San grazing lands, direct conflicts and raids of their life stock by Afrikaner frontier farmers gradually forced them out of the Western Cape (Elphick and Malherbe 1989). The small pox epidemic of 1713 was particularly harsh on the Khoe-San population with only 1 in 10 surviving. After the epidemic there were very few Khoe-San individuals left in the Western Cape (Elphick and Malherbe 1989). Continual wars against the Khoe-San, systematic killing of the Khoe-San, and capturing of Khoe-San women and children as so-called "inboekelinge" (to force them into a serfdom/indentured labour system) continued to reduce their numbers in areas occupied by Afrikaner frontier farmers (Elphick and Malherbe 1989; Shell 1994; Terreblanche 2002).

*2. European colonists*

Europeans started independent settlement at the Cape from 1658. Since they were to some extent independent of the Dutch East Indian Company (DEIC) they were initially called "free citizens" and as the populations' identity evolved they were referred to as Afrikaners (although "Africander" was initially used to indicate locally born slaves). The population had a high growth rate of about 2.6% per annum (Ross 1993). Although a small part of this increase was the result of a continuous arrival of immigrants, it is mostly due to the population's high fecundity (Figure S12).

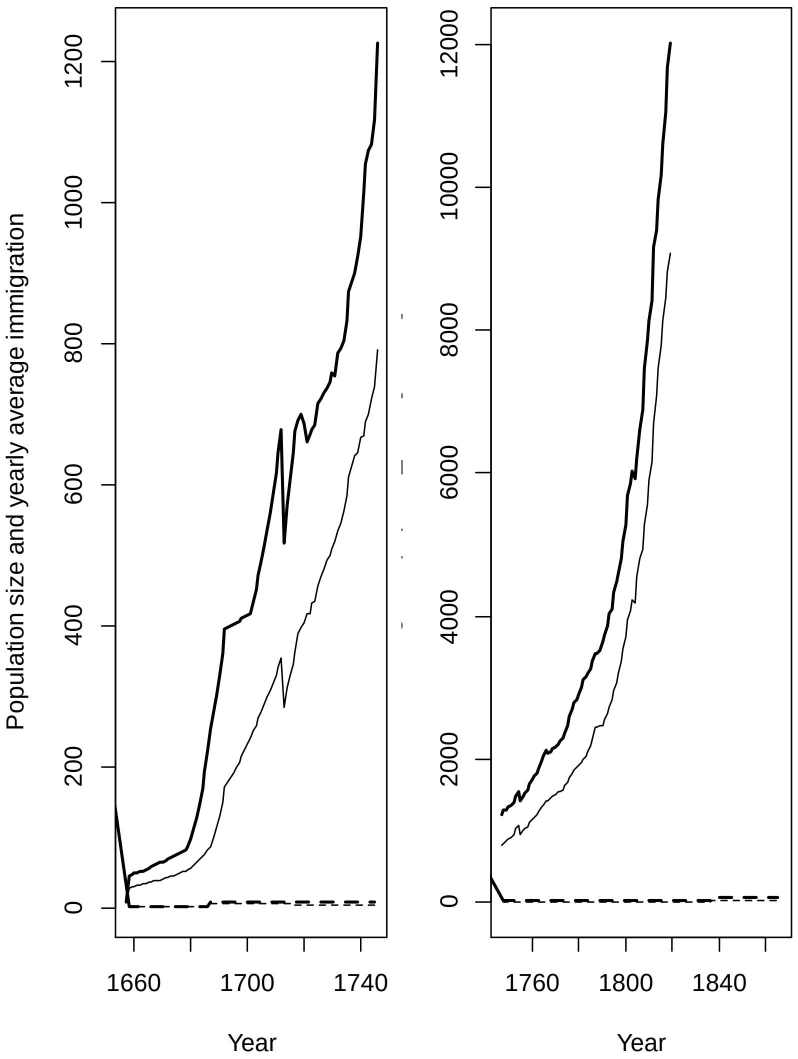


Figure S12. The population size of adult Afrikaner men (solid heavy line) and women (solid lighter line) as a function of time. Data was obtained from Gouws (1981). The yearly average number of immigrants that married into Afrikaner families is given in dashed lines with men in heavy and women in lighter lines. Data was counted from Heese (1971). These values are averages for 30 year periods, so for instance, from 1808 to 1837, 970 men who were not born at the Cape, got married in the colony, giving a yearly rate of 32⅓. Note the different scales on the y-axes. The Y-axis in the second panel starts where the axis ends in the first panel.

*3. Slaves*

Starting with two boats carrying about 400 slaves from West Africa in 1658 the slave population grew to 36,169 in 1834 when slavery was abolished at the Cape (Armstrong & Worden 1989). The total number of slaves and their origin has to be inferred indirectly, because records are missing and many arrivals were not recorded (Worden pers comm). Even so, Shell (1994) argued that about 63,000 slaves arrived in the colony between 1658-1807. He calculated that roughly a quarter came from each of; Africa (26.4%, east coast, excepting first two shiploads), Madagascar and

Mascarenes (25.1%), South Asia (25.9%) and Southeast Asia (22.7%) (Shell 1994). Based on household inventories, Worden (2016a 2016b) estimated that more slaves came from Asia, especially South Asia, and that substantially fewer came from Madagascar and Africa. Both sources suggest marked temporal variation in the origin of slaves, with Asian slaves predominating in the 17th and earlier 18th centuries and African slaves dominating towards 1807 (Figure S13). Initially the DEIC owned most slaves but after 1692 colonists owned the majority (Armstrong and Worden 1989). Shortly after 1710 slaves outnumbered the colonists and this situation remained until the abolition of slave trade in 1807 (Armstrong and Worden 1989). Although the slave population creolized (>50% locally born) in 1770 (Shell 1994), the local growth was lower than that of the colonists (Armstrong and Warden 1989). After emancipation in 1834 slaves were freed after a four-year apprenticeship and formed the backbone of the Coloured community.

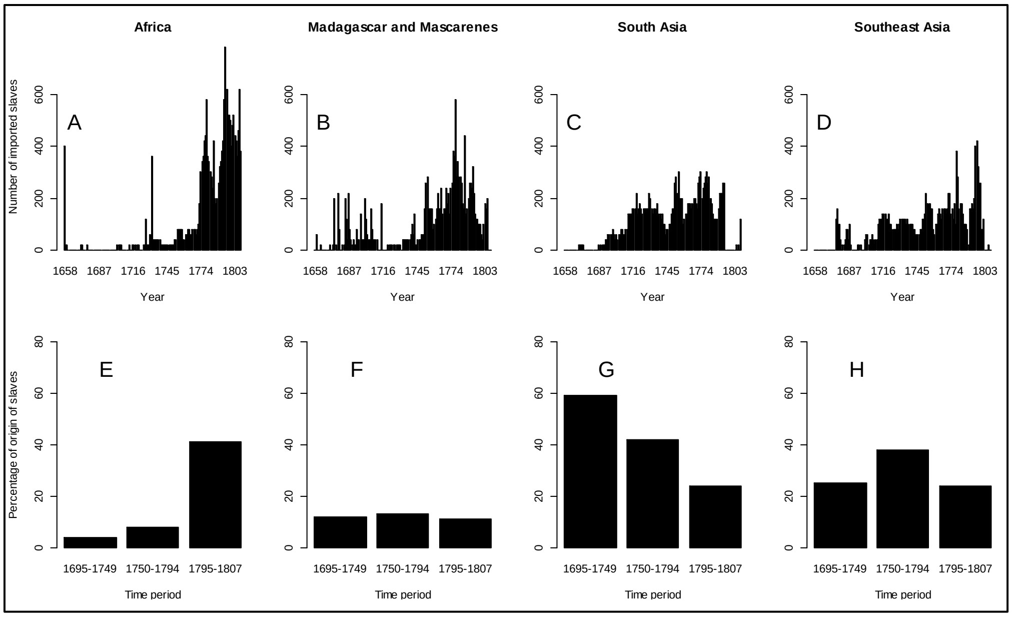


Figure S13. The origins of slaves arriving in the Cape. Estimates based on Shell (1994; A-D) and Worden (2016, workshop; E-H). Shell's (1994) data were read off his figure 2-1, rounded to the nearest 20, with slight adjustments to 1) fit the time line of imports from 1658 – 1807, and to 2) correct for the first two vessels that arrived from Western Africa in 1658. Worden's (2016b) estimates come from his Appendix. Slaves were from Africa (A & E), Madagascar and Mascarenes (B & F), South Asia (C and G) and Southeast Asia (D & H).

**Genealogical evidence of admixture**

Historians in South Africa compiled genealogical registries for the Afrikaner population based on church records (summarized in Greeff & Erasmus (2015)). From these records it is clear that the Afrikaner population is admixed (Table S5). The estimates vary, but the non-European contribution is between 5.4 and 7.2 percent.

People that are given the toponym "van die Kaap" (meaning from the Cape) rather than a surname are of uncertain heritage. While these individuals were born at the Cape their parent(s) may have been a slave, Khoe-San or even European (Ball undated). For this reason and for other reasons given in the main text, a genetic analysis is required to make an accurate calculation of the population's constitution.

Table S5. Four estimates from three studies of the percentage composition of Afrikaners.

| Origin of contributions | All immigrants up to 1867 weighted by entry time[a] | All immigrants up to 1807 weighted by entry time[a] | 1063 baptisms in 1807 in Cape[b] | One living individual[c] |
|---|---|---|---|---|
| **European** | **89.6[d]** | **89.3** | **90.7** | **93.34** |
| Netherlands | 34.8 | 36.8 | 34.1 | 37.22 |
| German | 33.7 | 35.0 | 29.2 | 27.21 |
| French | 13.2 | 14.6 | 24.7 | 26.26 |
| Other European | 7.9 | 2.9 | 2.7 | 2.65 |
| **Non-Euroean** | **6.9** | **7.2** | **5.4** | **6.04** |
| East-Asian | | | | 1.89 |
| South-East Asian | | | | 0 |
| South Asian | | | | 1.66 |
| India | | | | 1.66 |
| Africa | | | | 2.48 |
| Madagaskar | | | | 0.06 |
| Guinea | | | | 0.27 |
| Van die Kaap[e] | | | | 2.15 |
| **Unknown** | **3.5** | **3.5** | **3.9** | **0.62** |

[a] Heese (1971) divided 1657-1867 into seven 30-year slots (and for the period 1657-1807 in 5 30-year slots), for each slot new immigrants are listed and their contribution was weighted by their number of children and each earlier time-slot was weighed twice as heavily as the subsequent time-slot. Hence, an immigrant in 1658 contributed 64 "blood units" per child whereas an individual in 1840 contributed 1 per child.
[b] de Bruyn (1976) quantified the average composition of 1063 children baptized in the Cape in 1807. This sample represents about 15% of the entire Afrikaner population at the time.
[c] Percentages divided by 82.81 from Greeff (2007). It corrects for incomplete lineages.
[d] numbers in bold are totals of normal type indented from each area listed underneath.
[e] The expression "van die Kaap" was given as a toponymical surname and means "from the Cape". It was given to children born at the Cape and was presumably used to hide the fact that their parents were slaves or manumitted slaves. However, their parents may have been Khoe-San or even European (Ball, 2006).

**Genetic structure of the Afrikaner population**

Surprisingly little is known about the genetic structure of the Afrikaner population. The few genetic studies on Afrikaners suggested admixture between European and non-European populations. 1) For some loci Afrikaners have more alleles than Europeans (Erasmus et al. 2015) and 2) Afrikaners have non-European alleles at some loci (Nurse et al. 1985). Botha & Pritchard

(1972) estimated that slaves from Africa and Asia, and/or Khoe-San, contributed between 6 – 7% of genes to the Afrikaner population. Botha and Pritchard (1972) based their estimate on blood group allele frequencies.

A number of familial diseases are more common in Afrikaners than in European populations (Dean, 1963; Cilliers & Beighton, 1990; Jenkins et al. 1980; Hayden et al. 1980; Rosendorff et al. 1987; Torrington & Viljoen 1991; Torrington et al. 1986; Heyl, 1970; Beighton et al. 1977). These data supports the notion of a significant founder effect during the formation of the population (Botha & Beighton, 1983a, b; Nurse et al. 1985). While a deep routing pedigree of one Afrikaner clearly illustrated the existence of many common ancestors, these were so distant that his pedigree-inbreeding coefficient was not higher than European averages (Greeff 2007).
